# Cardiomyocyte-fibroblast interaction regulates ferroptosis and fibrosis after myocardial injury

**DOI:** 10.1101/2023.02.07.527364

**Authors:** Mary E. Mohr, Shuang Li, Allison M. Trouten, Rebecca A. Stairley, Patrick L. Roddy, Chun Liu, Min Zhang, Henry M. Sucov, Ge Tao

## Abstract

Neonatal mouse hearts have transient renewal capacity which is lost in juvenile and adult hearts. After myocardial infarction (MI) in neonatal hearts, an initial loss of cardiomyocytes occurs but it is unclear through which type of regulated cell death (RCD). In the current studies, we induced MI in neonatal and juvenile mouse hearts, and show that ischemic cardiomyocytes primarily undergo ferroptosis, a non-apoptotic and iron-dependent form of RCD. We demonstrate that cardiac fibroblasts (CFs) protect cardiomyocytes from ferroptosis through paracrine factors and direct cell-cell interaction. CFs show strong resistance to ferroptosis due to high ferritin expression. Meanwhile, the fibrogenic role of CFs, typically considered detrimental to heart function, is negatively regulated by paired-like homeodomain 2 (Pitx2) signaling from cardiomyocytes. In addition, Pitx2 prevents ferroptosis in cardiomyocytes by regulating ferroptotic genes. Understanding the regulatory mechanisms of cardiomyocyte survival and death can identify potentially translatable therapeutic strategies for MI.

## INTRODUCTION

Heart failure caused by MI is one of the main causes of death worldwide ^1^. Following MI, increased reactive oxygen species (ROS) promotes cardiomyocyte death and maladaptive fibrosis ^2, 3^. Tightly regulated cell death and cellular injury response are crucial for tissue repair and maintenance of cardiac function. It is unclear to what extent each type of RCDs contributes to the loss of cardiomyocytes in the infarcted myocardium. Widely studied RCDs in heart disease includes apoptosis, necroptosis, and autophagy ^4–6^. Ferroptosis, a recently added RCD member, is a non-apoptotic form of RCD caused by excessive production of lipid hydroperoxides in the presence of iron ^7^. Ferroptotic cells do not demonstrate apoptotic hallmarks such as caspase expression or plasma membrane rupture ^7^. Morphologically, ferroptosis is characterized by shrinkage of mitochondria and mitochondrial membrane rupture ^8, 9^. Cellular iron, normally stored in a protein complex composed of ferritins, is required for the onset of ferroptosis ^10^. Strong rationale for studying ferroptosis in the heart includes the known cardiotoxicity of iron and the presence of iron overload in the border zone of infarcted myocardium ^11^. Two enzymatic proteins have been identified as the major inhibitors of ferroptosis. The glutathione peroxidase 4 (Gpx4) inhibits ferroptosis by reducing lipid peroxides to lipid alcohols using its main cofactor glutathione (GSH) ^12^. Meanwhile, the ferroptosis suppressor protein 1 (Fsp1) functions as an oxidoreductase on cell membrane to reduce coenzyme Q10 (CoQ) and generates antioxidants to halt the propagation of lipid peroxides ^13^, ^14^. Recently, increased ferroptosis was observed after ischemic injury and cardiomyopathy in adult mouse models ^15^, ^16^.Little is known about the transcriptional regulation of ferroptosis in cardiomyocytes and how ferroptotic cells affect surrounding cells after myocardial injury.

In cancer cell models, high cell density in culture prevents ferroptosis ^17^. In infarcted myocardium, cell density decreases with the loss of cardiomyocytes, and is restored when resident fibroblasts are activated and migrate to the infarct zone ^18^. CFs are fast responders to cardiac injury. They lay down extracellular matrix (ECM) to form fibrotic scarring, which is vital for the maintenance of myocardial integrity after heart attack ^19^. Cardiomyocyte-fibroblast interaction plays a role in maintaining cardiac function ^20^, ^21^, although how this cell-cell interaction impacts cardiomyocyte death remains unanswered.

Neonatal mammalian hearts are regenerative due to residual cell cycle activity in cardiomyocytes that is lost in juvenile and adult hearts ^22^, ^23^. Utilizing mouse heart surgery models and *in vitro* human cardiac cell cultures, we aimed to address whether and how ferroptosis was involved in the regeneration of young hearts after injury. We found that ferroptosis contributes significantly to the loss of cardiomyocytes after MI in postnatal day (P) 1 and P7 hearts. CFs protect cardiomyocytes from ferroptosis through cell-cell interaction and paracrine signaling. CFs are resistant to ferroptosis and can tolerate high iron load. Mechanistically, the Pitx2 signaling in cardiomyocyte prevents ferroptosis and negatively regulates the fibrotic activity of CFs.

## RESULTS

To analyze the prevalence of common types of RCDs in cardiomyocytes during MI, we performed left anterior descending coronary artery occlusion (LAD-O) in the regenerative neonatal mouse heart at P1 and the non-regenerative P7 hearts ^24^. Immunofluorescence staining of tissue from the infarcted left ventricular free wall showed a low rate of cardiomyocytes positive for the apoptotic cleaved Caspase 3 (cCasp3) at 1, 3, and 6 day-post-MI (DPMI) after P1 or P7 LAD-O (Figure 1A-G). The ratio of necroptotic cardiomyocytes, marked by phosphorylated mixed lineage kinase domain like pseudokinase (pMLKL) ^25^, were below 1% at all time points (Figure 1H-N). However, more pMLKL-positive cardiomyocytes were observed at 6 hours after P1 or P7 LAD-O, likely caused by acute ischemic injury (Figure S1A-D). Next, we examined the level of prostaglandin-endoperoxide synthase 2 (Ptgs2), a commonly used ferroptotic marker (Figure 1O-U) ^12^. Ptgs2 was detected in 7% of cardiomyocytes at 1 DPMI after P1 LAD-O, and the ratio dropped at 3 DPMI and decreased further at 6 DPMI (Figure 1O-Q, U). In comparison, after P7 LAD-O, around 10% of ferroptotic cardiomyocytes were detected in all time points, indicating a consistent loss of cardiomyocytes caused by ferroptosis (Figure 1R-U). Markers of these three types of RCD were barely detectable in cardiomyocytes after sham-operated procedure (data not shown). These data indicate that ferroptosis is the major form of RCD in both P1 and P7 hearts after LAD-O.

**Figure 1.**
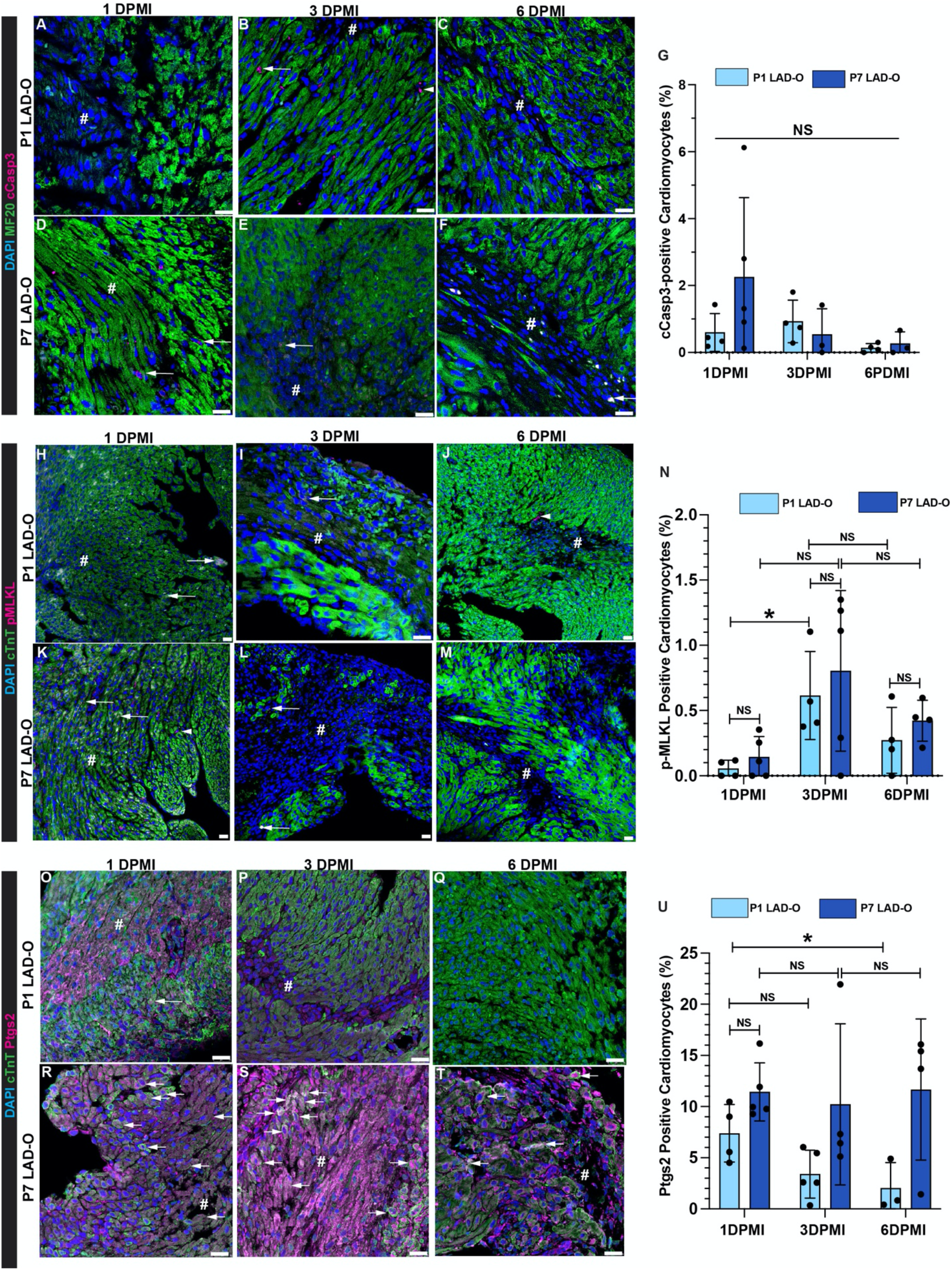
Ferroptosis is more prevalent in cardiomyocyte than apoptosis and necroptosis in infarcted postnatal hearts. LAD-O was performed at either P1 or P7, heart tissue section prepared at 1, 3 or 6 DPMI. (A-G) Infarct zone stained for cCasp3 (Magenta), MF20 (green) and DAPI (blue). Arrow, cardiomyocytes positive for cCasp3, ratio quantified in (G). Arrowhead, non-cardiomyocyte positive for cCasp3. (H-N) Infarct zone stained for pMLKL (Magenta), cTnT (green) and DAPI (blue). Arrow, cardiomyocytes positive for pMLKL, ratio quantified in (N). Arrowhead, non-cardiomyocyte positive for pMLKL. (O-U) Infarct zone stained for Ptgs2 (Magenta), cTnT (green) and DAPI (blue). Arrow, cardiomyocytes positive for Ptgs2, ratio quantified in (U). Error bars indicate SD. *, p<0.05. NS, not significant. NS in A, no significance among all groups. #, infarct zone. Scale bar, 25 μm (A-F, H-M, O-T). See also Figure S1 and S2.

Ferroptosis is comparatively new to the family of RCD, and the identification of reliable protein markers has not reached a consensus. Therefore, we confirmed our findings by immunofluorescence staining of 4-Hydroxynonenal (4-HNE), a byproduct of lipid peroxidation, considered the primary cause of ferroptosis (Figure S2A-C) ^15^. The level of 4-HNE is detectable in cardiomyocytes after P1 LAD-O, but at a significantly lower rate compared to P7 LAD-O (Figure S2A-C). At the regenerative stage, MI does lead to increased level of lipid peroxidation as confirmed by malondialdehyde (MDA) assay (Figure S2D). Noticeably, although LAD-O at P7 caused higher 4-HNE level at 1 and 3 DPMI, it dropped to comparable level at 6 DPMI (Figure S2C). However, 4-HNE is still present in cardiomyocytes at 21 days after P7 LAD-O, suggesting a persistent oxidative stress in the non-regenerative myocardium after injury (Figure S2E, F). Ptgs2 is not detectable at this point (Figure S2G), indicating that the infarct is stabilized after remodeling and loss of cardiomyocytes is low.

The ferroptotic pathway has become a popular target for anti-tumor research, and it has been reported that high density/confluency protects cells from ferroptosis ^17^. Cardiomyocytes are packed in the myocardium at a high density which decreases after MI. We hypothesized that the influx and expansion of CFs can restore cardiac cell density and therefore protect the remaining cardiomyocytes from ferroptosis. To test this possibility, human induced pluripotent stem cells (hiPSCs) expressing GFP-tagged TITIN were differentiated into iPSC-derived cardiomyocytes (iCMs) and cultured at low, mid, or high density (Figure S3A) ^26^. Cells were treated with erastin, a ferroptosis-inducing System Xc- inhibitor ^27^. Erastin caused comparable death rate in iCMs cultured at different cell densities (Figure 2A). On the contrary, cultured primary human cardiac fibroblasts (HCFs) had significantly higher resistance to erastin at mid and high density, compared to low density (Figure 2B). Next, we tested if iCMs can be protected from erastin when total cell density is increased by co-seeding HCFs. iCMs were either cultured alone, or co-cultured to confluency with HCFs (Figure 2C-F). Co-culture with HCFs significantly decreased the level of PTGS2 in iCMs after erastin treatment (Figure 2C-G), suggesting a protective effect caused by HCFs. It is noteworthy that while the iCMs are sensitive to erastin even at high culture density, a human cardiomyocyte cell line called AC16 showed density-dependent resistance to erastin (Figure S3B, C) ^28^. Compared to the iCMs which beat spontaneously and do not proliferate efficiently, the immortalized AC16 cells do not beat and are highly proliferative ^28^. The AC16 cells do not have organized sarcomere structure and resemble the morphology of embryonic cardiomyocytes (data not shown). These characterizes of AC16 cells suggest certain level of dedifferentiation which could explain the different response to erastin treatment.

**Figure 2.**
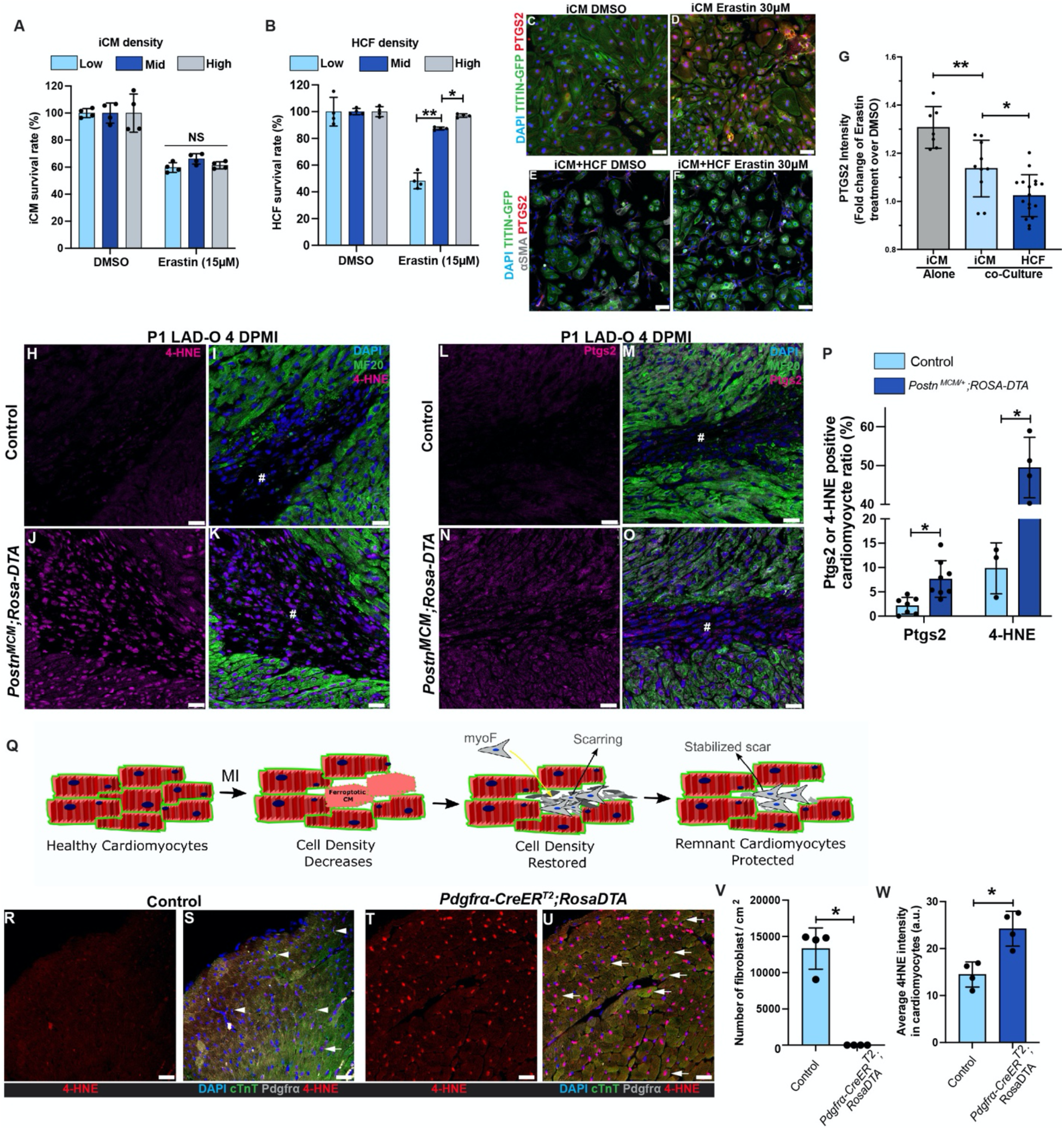
Cardiac fibroblasts protect cardiomyocytes from ferroptosis. (A) Survival (negative for trypan blue) rates of iCMs cultured at low, mid or high density after erastin treatment (15 μM, 5 hours). (B) Survival rates of HCFs cultured at low, mid or high density after erastin treatment (15 μM, 5 hours). (C, D) PTGS2 (red) and DAPI (blue) were stained and imaged with endogenous TITIN-GFP (green) in iCM after erastin (30 μM) treatment. (E, F) PTGS2 (red), aSMA (grey) and DAPI (blue) were stained and imaged with endogenous TITIN-GFP (green) in co-cultured iCMs and HCFs after erastin (30 μM) treatment. (G) Fold change of PTGS2 fluorescent intensity in iCMs and HCFs. (H-O) Heart tissue of controls (*Postn^MCM/+^*, H, I, L, M) and *Postn^MCM/+^;ROSA-DTA* (J, K, N, O) mice were stained for 4-HNE (magenta, H-K) or Ptgs2 (magenta, L-O) at 4 DPMI after P1 LAD-O, tamoxifen was administrated daily at 1-3 DPMI. Green, MF20; blue, DAPI. #, infarct zone. (P) Ratio of cardiomyocytes positive for Ptgs2 or 4-HNE. (Q) Schematic of cardiac fibroblasts entering scar zone to increase cardiac cell density and rescue cardiomyocyte from ferroptosis. (R-U) Heart sections of 2-month-old control (*ROSA-DTA*, R, S) and *Pdgfra-CreER^T2^;ROSA-DTA* (T, U) mice stained for 4-HNE (red), cTnT (green), Pdgfra (grey) and DAPI (blue) after tamoxifen administration (see Figure S3). (V) Density of Pdgfra-positive cells in control and mutant groups. (W) Average 4-HNE intensity in cardiomyocytes from both groups. #, infarct zone. Error bars indicate SD. *, p<0.05. **, p<0.01. NS, not significant. NS in A, no significance among three groups. Scale bar, 50 μm (C-F), 25 μm (H-O, R-U). See also Figure S3.

We next determined if CFs protect cardiomyocytes from ferroptosis *in vivo*. We crossbred *Postn^MCM/+^* mice, a tamoxifen-inducible myofibroblast-specific *Cre* strain, to *Rosa-DTA* strain ^19^. Upon tamoxifen injection, Cre-induced diphtheria toxin A subunit (DTA) expression eliminates the myofibroblasts with Periostin (*Postn*) gene activity (Figure S3D-G). Myofibroblast, known as activated fibroblasts, are essential for myocardial remodeling after MI and can be identified by alpha smooth muscle actin (aSMA), Vimentin (Vim), and Postn ^18^. *Postn^MCM/+^;Rosa-DTA* mice had significantly higher level of 4-HNE (Figure 2H-K) and Ptgs2 (Figure 2L-O) in the infarct and border zone compared to the control group (Figure 2P). These results suggest that, after the loss of cardiomyocytes caused by ferroptosis in infarcted myocardium, the influx of CFs can restore the total cell density and protect remaining cardiomyocytes from further ferroptosis (Figure 2Q).

We further examined if CFs in the adult mouse heart protect cardiomyocytes. Loss of myofibroblasts after MI compromises cardiac muscle integrity and affects myocardial remodeling ^19^, ^29^. This may cause a secondary defect in cardiomyocytes that obstruct our investigation on the roles of CFs. To rule out this possibility, we bred *Pdgfrα-CreER^T2^;Rosa-DTA* adult mice and administered tamoxifen to eliminate cardiac interstitial fibroblasts from healthy myocardium without MI (Figure S3H) ^30^. To avoid potential secondary effects, our strategy was optimized so that the myocardial interstitial fibrosis and contractile functions were not affected (Figure S3I-K).

With effective removal of the myocardial mesenchymal cell population, the *Pdgfrα-CreER^T2^;Rosa-DTA* hearts had significantly higher 4-HNE level in cardiomyocytes (Figure 2R-W), thus supporting the hypothesis that cardiac fibroblasts directly regulate cardiomyocyte homeostasis.

To investigate the mechanisms underlying the resistance of CFs to ferroptosis (Figure 2B), we started with characterizing and authenticating the primary HCFs used in current studies. In culture, HCFs demonstrate high level of αSMA and TGFβR2, and nucleus-localized pSMAD2 and pSMAD3 (Figure 3A-C). These characteristics of myofibroblasts make the HCFs appropriate for the current studies. Cardiomyocytes and CFs were exposed to increased ROS after myocardial injury ^31^. *In vitro*, H_2_O_2_ treatment reduced survival rate of both iCMs and HCFs on a dosage-dependent pattern indicating their sensitivity to oxidative stress (Figure 3D, Figure S3L) ^31^. When treated with erastin at low to high dosages, HCFs survival rate decreased with dosage-dependency from 2 μM to 10 μM but recovered when erastin concentration reached 20 μM (Figure 3E, F). When erastin reached 30 μM, HCF survival rate bounced back to be comparable to 2uM (Figure 3F). The same resistance to high erastin dosage was not observed in iCMs (Figure S3M). To validate the observed resistance of HCFs to ferroptosis in mouse models, we prepared primary cultures of CFs from P1 and P7 mouse heart ventricles ^32^. A similar pattern of resistance was observed in P1 CFs, with the survival rate decreasing and subsequently increasing as the erastin dosage increased (Figure 3G). The P7 group also showed an increased but milder resistance compared to the P1 group (Figure 3H). In comparison, HEK-293, a human embryonic kidney epithelial cell line, had decreased survival in a dosage-dependent manner when treated with either H_2_O_2_ or erastin (Figure 3I, J). High confluency did not protect the HEK-293 cells from erastin-induced ferroptosis (Figure 3K). It is likely that the ferroptosis-resistance is not ubiquitous but restricted to certain cell, injury and/or stress types.

**Figure 3.**
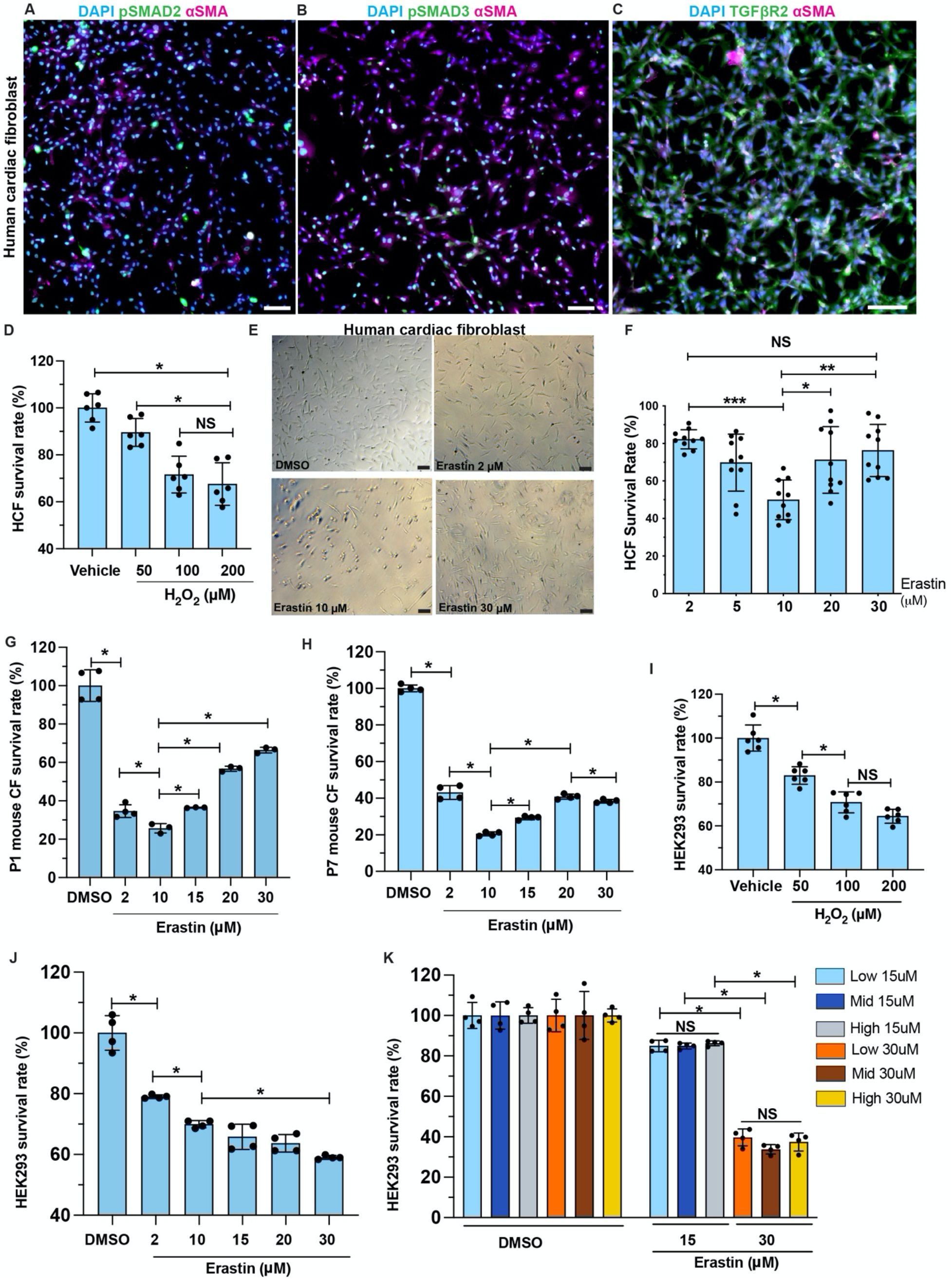
Cardiac fibroblasts are resistant to ferroptosis. (A-C) HCFs were stained for pSMAD2 (green in A), pSMAD3 (green in B) and TGFβR2 (green in C), with aSMA (magenta) and DAPI (blue). (D) Survival rate of HCFs after H_2_O_2_ or vehicle (H_2_O) treatment. (E, F) Brightfield of HCFs after erastin or DMSO treatment. Survival rate quantified in F. (G, H) Survival rate of primary mouse cardiac fibroblasts (CF), prepared from P1 (G) and P7 (H) hearts, after erastin treatment. (I) Survival rate of HEK293 cells after H_2_O_2_ or vehicle treatment. (J) Survival rate of HEK293 cells treated with erastin at gradient concentration. (K) Survival rate of HEK293 cells cultured at low, mid, and high density after erastin treatment at 15 or 30 μM, compared to DMSO groups. Error bars indicate SD. *, p<0.05; **, p<0.01; ***, p<0.001. NS, not significant. Scale bar, 100 μm (A-C, E). See also Figure S3.

Next, we aimed to determine the cellular and molecular mechanisms underlying the protective roles of CFs. Considering the proximity between cardiomyocytes and CFs in myocardium, we hypothesized a paracrine-mediated and a contact-directed mechanisms. HCFs were pretreated with H_2_O_2_ to mimic the oxidative stress in infarcted myocardium, then H_2_O_2_ was removed, and HCF-conditioned medium was prepared (Figure 4A). Conditioned media from H_2_O_2_-pretreated HCFs, but not vehicle-treated cells, partially rescued iCMs from erastin-induced ferroptosis, compared to non-conditioned medium (Figure 4B). We then used a protein array assay to examine a pool of common paracrine cytokines and chemokines. H_2_O_2_ treatment significantly increase the secretion of Interleukin-8 (IL-8) and showed a trending increment of Epidermal Growth Factor (EGF) level (Figure 4C, D). IL-8, also known as CXCL8, is typically released by macrophages and epithelium to attract neutrophils during inflammation ^33^. Meanwhile EGF family ligands such as Neuregulin are well known for their key roles in cell proliferation, including in cardiomyocyte ^34^. Treating iCM with either IL-8 or EGF recombinant protein promoted iCM survival after erastin treatment (Figure 4E, F). These results suggest a stoichiometric effect of a series of paracrine factors from CFs that contribute to cardiomyocyte survival after MI. One remaining question is how cardiomyocytes respond to IL-8 efficiently. Cxcr1 and Cxcr2 are known receptors of IL-8 ^35^. To examine whether *Cxcr1* and *Cxcr2* are expressed in regenerating neonatal cardiomyocytes, we prepared RNA from purified cardiomyocyte nuclei at 4 DPMI after P1 LAD-O. qPCR showed similar level of *Cxcr1* in cardiomyocytes compared to total blood cell RNA (Figure 4G). *Cxcr2* level is lower in cardiomyocyte but still detectable (Figure 4G). These findings support the hypothesis that fibroblasts confer protection on cardiomyocytes through paracrine signaling.

**Figure 4.**
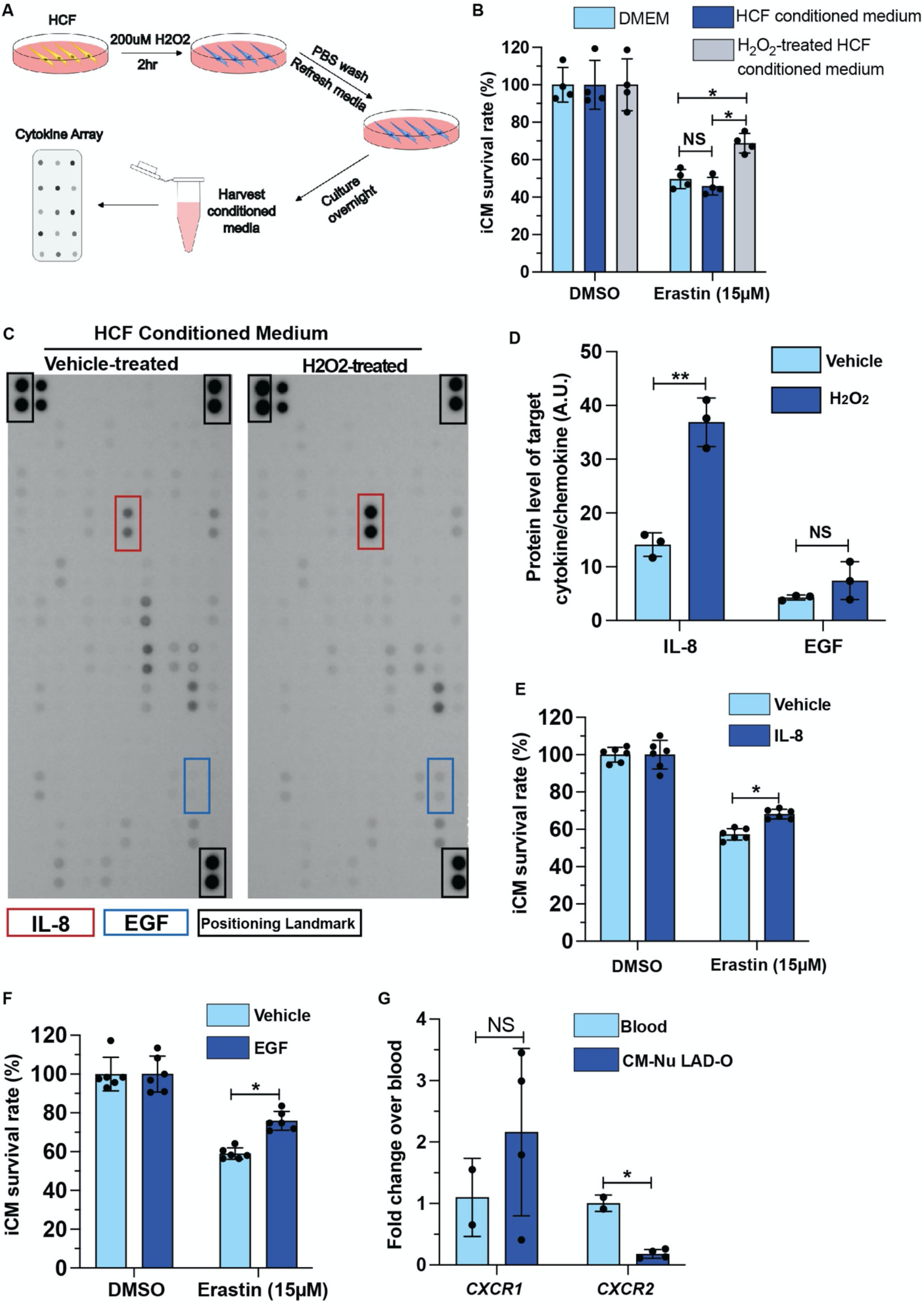
Fibroblast-derived cytokine and chemokine promote iCM survival after erastin treatment. (A) Flowchart of conditioned media and cytokine array experiment. (B) Survival rate of iCMs, cultured in control medium (DMEM), HCF conditioned medium, or H_2_O_2_-treated HCF conditioned medium, after erastin treatment, normalized to respective DMSO control groups. (C) Cytokine-chemokine protein array blotting image of conditioned media from vehicle- or H_2_O_2_-treated HCFs. (D) Blotting signal intensity of IL-8 and EGF. (E, F) Survival rate of iCMs after erastin treatment in the presence of IL-8 (E) or EGF (F), normalized to DMSO control groups. (G) qPCR of *CXCR1* and *CXCR2* in total blood cells and purified cardiomyocyte nuclei after P1 LAD-O. Error bars indicate SD. *, p<0.05; **, p<0.01. NS, not significant.

In terms of the contact-directed mechanism for CFs to rescue ferroptotic cardiomyocytes, we had a hint from the immunofluorescence staining of ferritin heavy chain 1 (Fth1) and ferritin light chain (Ftl). Known as the key components of the anti-ferroptotic and iron-storing ferritins, Ftl and Fth1 form protein cages while Fth1 oxidizes ferrous iron (Fe^2+^) to ferric iron (Fe^3+^), which is then stably stored in the mineral core ^10, 36, 37^. At one day after P7 LAD-O, a group of Fth1-high cardiac cells lined the border of ischemic (cTnT-low) and healthy (cTnT-high) myocardium (Figure 5A). A similar pattern was observed when staining Ftl (Figure 5B). Co-staining suggests these cells are a subgroup of Pdgfrα-expressing fibroblasts (Figure 5C, D). The presence of this subpopulation of fibroblasts persists into the later period of heart regeneration (Figure 5E, F), and were also observed after P1 LAD-O (Figure S4A, B). Thus, we further hypothesized that these ferritin-high fibroblasts establish cell-cell contact and share the iron-burden with cardiomyocytes at the border zone (Figure 5G) ^11^. The presence of Cx45 and Cx43, known to form gap junctions between cardiomyocytes and fibroblasts ^38^, indicated direct cell-cell contact between these cardiac cells (Figure 5H, I). Importantly, gap junctions have an internal diameter large enough for shuttling free iron ^39^. *In vivo*, cardiac fibroblasts showed increased accessibility at *Gja1* (encoding Cx43) and *Gjc1* loci (encoding Cx45) after MI (Figure S4C). In HCFs, CX43 and CX45 are expressed and conjugate at the perinuclear area when the cell density is low but distribute more to the cytoplasm and plasma membrane when cell density is high (Figure S4D-G’). Co-culture of iCMs and HCFs showed the presence of CX43 and CX45 between the two cell types (Figure S4H, I).

**Figure 5.**
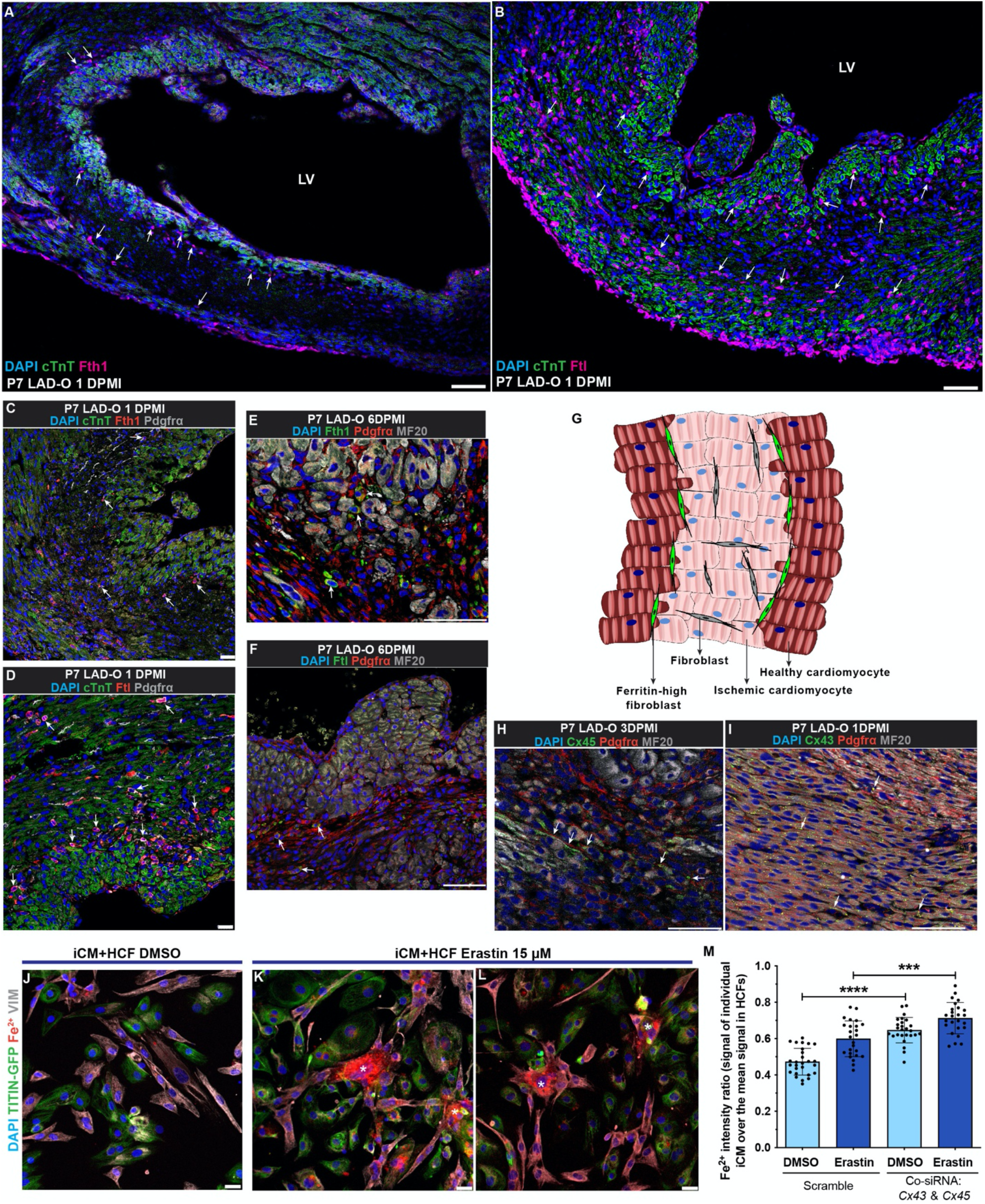
Cardiac fibroblasts interact with cardiomyocytes through gap junctions to share free iron. (A, B) Wild type mouse heart tissue stained for Fth1 (magenta, A) or Ftl (magenta, B), with cTnT (green) and DAPI (blue) at 1DPMI after P7 LAD-O. Arrows, non-cardiomyocytes positive for Fth1 (A) or Ftl (B). (C, D) Mouse heart tissue stained for Fth1 (red, C) or Ftl (red, D), with Pdgfrα (grey), cTnT (green) and DAPI (blue) at 1 DPMI after P7 LAD-O. Arrows, cells positive for Pdgfrα and Fth1 (C) or Ftl (D). (E, F) Mouse heart tissue stained for Fth1 (green, E) or Ftl (green, F), with Pdgfrα (red), MF20 (grey) and DAPI (blue) at 6 DPMI after P7 LAD-O. Arrows, cells positive for Pdgfrα and Fth1 (E) or Ftl (F). (G) Diagram of cardiomyocyte-fibroblast interaction after MI. (H, I) Mouse heart section stained for Cx45 (green, H) or Cx43 (green, I) with Pdgfrα (red), MF20 (grey) and DAPI (blue) after P7 LAD-O. Arrows, potential locations of gap junctions between cardiomyocytes and fibroblasts. (J-L) Co-cultured iCM and HCF stained for VIMENTIN (VIM, Grey), free Fe^2+^ (red), DAPI (blue), and imaged with TITIN-GFP (green) after DMSO (J) or erastin (15 μM) (K, L) treatment. Asterisks, HCFs with accumulation of Fe^2+^. (M) siRNA knockdown of *CX43* and *CX45* simultaneously in iCM-HCF co-culture, Fe^2+^ fluorescent intensity ratio of iCMs over HCFs was quantified after erastin or DMSO treatment. LV, left ventricle. Error bars indicate SD. ***, p<0.001; ****, p<0.0001. Scale bar, 75 μm (A, B, E, F, H, I), 25 μm (C, D, J-L). See also Figure S4 and S5.

Iron is required for the onset of ferroptosis and presents in the completed culture media ^7^. When cultured separately, iCMs had a 55% increase of free Fe^2+^ after erastin treatment, while the HCFs had only a 17% increase (Figure S4J-N). When co-cultured, Fe^2+^ accumulated in HCFs after erastin treatment, compared to adjacent iCMs (asterisks in Figure 5J-L). This shift of Fe^2+^ burden is likely mediated by gap junctions, as simultaneous knockdown of *CX43* and *CX45* led to increased free Fe^2+^ in iCMs (Figure 5M). These data support the hypothetical model of fibroblasts sharing iron burden with cardiomyocytes after MI. Moreover, the fact that HCFs can tolerate high iron load suggests a potential mechanism underlying their high resilience under ferroptotic stress. It is known that CF density increases in the infarct zone after MI. *In vitro*, we found out that increased cell density promotes FTH1 and FTL protein level in HCFs (Figure S5A-C). Efficient knockdown of either *FTH1* or *FTL* caused lower survival rate of HCFs when treated with 30 μM of erastin (Figure S5D-F), the dosage at which HCFs showed resistance (Figure 3F). These findings suggest a unique mechanism for CFs to promote cardiomyocyte survival in a pro-ferroptotic micro-environment.

So far, we described a beneficial role of CFs during myocardial regeneration. However, the detrimental effect of an overly large population of activated fibroblasts cannot be neglected. Compared to P7 LAD-O, the activation of CFs into myofibroblasts occurs slightly earlier after P1 LAD-O (Figure S6A-D). However, P7 LAD-O resulted in a bigger population of myofibroblasts at 6 DPMI, shown by fate-mapping using *Postn^MCM^;ROSA^mT/mG^* mice (Figure 6A-C).

**Figure 6.**
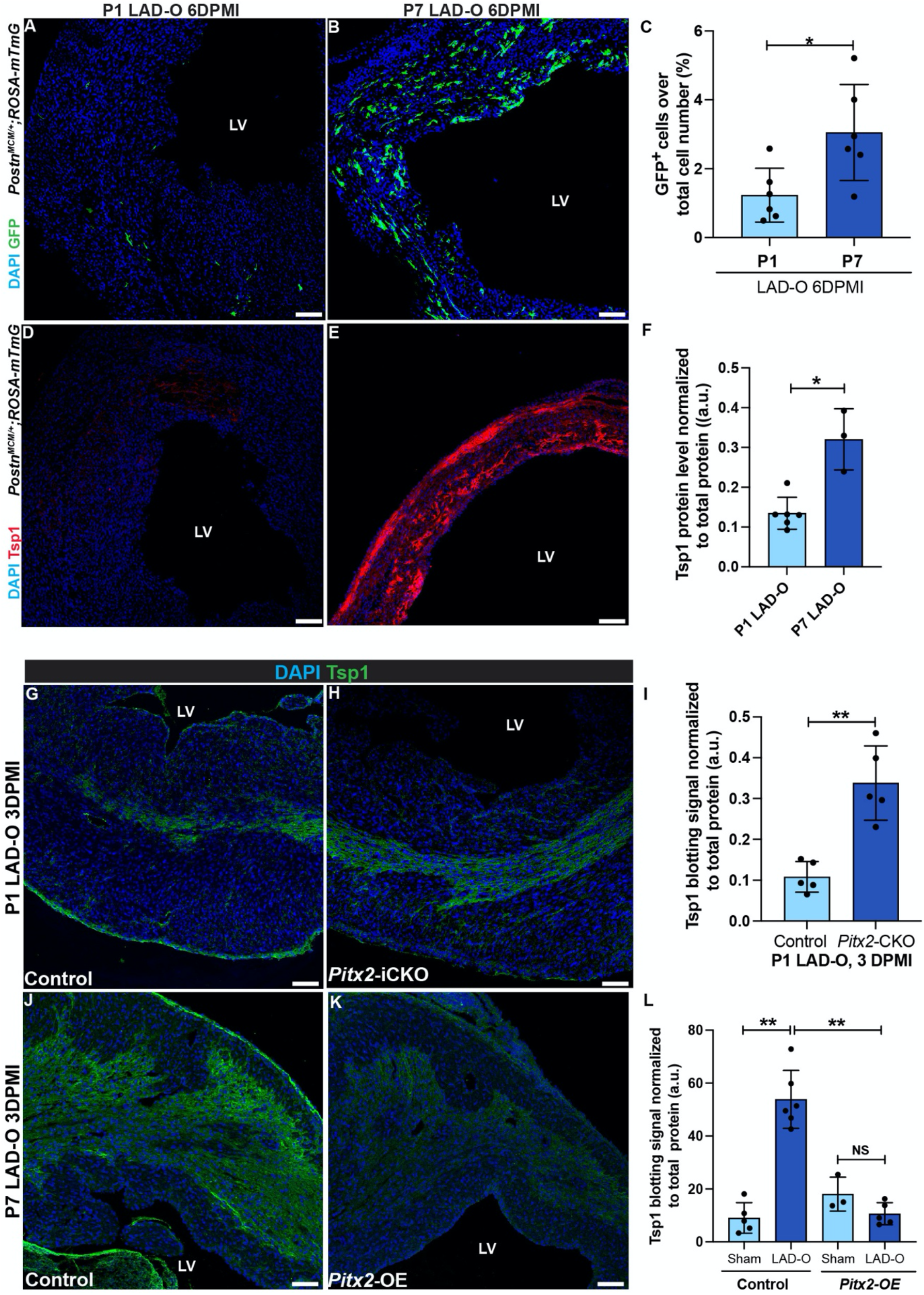
Pitx2 negatively regulates fibrosis by downregulating Tsp1 expression after MI. (A, B) *Postn^MCM/+^;ROSA-mTmG* heart sections imaged for endogenous GFP at 6 DPMI after P1 (A) or P7 (B) LAD-O. Tamoxifen administrated daily at 1, 2, 3 and 5 DPMI. (C) GFP-positive cell ratio over total nuclei number. (D, E) *Postn^MCM/+^;ROSA-mTmG* heart sections stained for Tsp1 (red) and DAPI (blue) at 6 DPMI after P1 (D) or P7 (E) LAD-O. (F) Normalized band (immunoblotting) intensity of myocardial Tsp1 at 4 DPMI after P1 or P7 LAD-O. (G, H) Control (*Mhc^cre-Ert^*) (G) and *Pitx2-iCKO (Mhc^cre-Ert^;Pitx2^f/f^*) (H) Heart tissue stained for Tsp1 (green) and DAPI (blue) at 3 DPMI after P1 LAD-O. Tamoxifen was given at P0-P2. (I) Normalized band intensity of Tsp1 in control and *Pitx2-iCKO* left ventricle at 3 DPMI after P1 LAD-O. (J, K) Control (*Mhc^cre-Ert^*) (J) and *Pitx2-OE (Mhc^cre-Ert^;Pitx2^gof^*) (K) Hearts were stained for Tsp1 (green) and DAPI (blue) at 3 DPMI after P7 LAD-O. Tamoxifen was given at P6-P8. (L) Normalized band intensity of Tsp1 in control and *Pitx2-OE* left ventricle at 3 DPMI after P7 LAD-O. Error bars indicate SD. *, p<0.05; **, p<0.01. NS, not significant. LV, left ventricle. Scale bar, 100 μm (A, B, D, E, G, H, J, K). See also Figure S6.

We examined key regulatory factors of fibroblast-to-myofibroblast transition using fluorescence staining and western blot. Thrombospondin 1 (Tsp1), encoded by *Thbs1*, is significantly higher at 6 DPMI after P7 LAD-O, compared to the P1 group (Figure 6D-F, Figure S6E). Tsp1 activates latent TGFβ1, which is essential for myofibroblast transition, collagen deposition and tissue scarring ^40^. We previously reported Pitx2 as a redox regulator that promotes neonatal mouse heart regeneration ^31^. Re-visiting earlier RNA-Seq data unveiled Pitx2 as a regulator of fibrotic pathways (Figure S6F-H)^31^. Cardiomyocyte-specific knockout of *Pitx2* upregulates the expression of a series of collagens and fibrotic regulators including *Thbs1* in injured myocardium (Figure S6F, G).

ChIP-Seq data showed *Thbs1* as a direct target of Pitx2, which binds to open chromatin region of *Thbs1* locus (Figure S6I). Tamoxifen-induced, cardiomyocyte-specific knockdown of *Pitx2 (Mhc^cre^-ERT2;Pitx2f/f, Pitx2*-iCKO) after P1 LAD-O led to increased Tsp1 in the infarct zone (Figure 6G-I, Figure S6J). On the contrary, overexpression of Pitx2 specifically in cardiomyocytes (*Mhc^cre-ERT2^;Pitx2^GOF^, Pitx2-OE*) reduced myocardial Tsp1 after P7 LAD-O (Figure 6J-L, Figure S6K). Together, these findings support a regulatory role of Pitx2 in fibrosis after MI.

Further analysis of RNA-Seq data generated in regenerating neonatal mouse hearts and iCMs indicated that Pitx2 is directly involved in a cell autonomous, anti-ferroptotic mechanisms ^31^. Pitx2 positively regulates a series of anti-ferroptotic genes including *Fth1, Ftl, glutathione synthetase* (*Gss*), CDGSH iron sulfur domain 1 (*Cisd1*) (Figure 7A, B) ^10^, ^12^, ^41^, as well as *Gpx4*, as reported previously ^31^. Pitx2 also represses lysophosphatidylcholine acyltransferase 3 (*Lpcat3*), required for the onset of ferroptosis (Figure 7A, B) ^42^. The expression of these Pitx2 target genes demonstrated an anti-ferroptotic pattern in control hearts after injury, compared to the control-sham hearts (Figure 7A). However, this anti-ferroptotic pattern was not noted in hearts with *Pitx2* knocked out specifically in the cardiomyocytes (Figure 7A). The regulation of ferroptotic genes by PITX2 was validated using qPCR in iCMs with erastin treatment and lentivirus overexpressing PITX2 (Figure 7C). Furthermore, our earlier Pitx2 ChIP-Seq showed the direct regulation of *Ftl, Fth1, Gpx4* and *Cisd1* by Pitx2 in regenerating neonatal mouse hearts (Figure 7D) ^31^. *In vitro, PITX2* transcript level increased in response to erastin treatment (Figure 7E), further supporting the direct involvement of PITX2 in the regulation of cardiomyocyte ferroptosis. In mouse models, immunofluorescence staining showed decreased 4-HNE in *Pitx2-OE* cardiomyocytes after LAD-O, compared to controls (Figure 7F, G). Western blot of left ventricular tissue from control and *Pitx2*-OE hearts showed no detectable Ptgs2 after sham procedure (Figure 7H, I). However, after LAD-O, *Pitx2-OE* left ventricles had lower level of Ptgs2 compared to controls, indicating lower rate of ferroptosis (Figure 7H, I).

**Figure 7.**
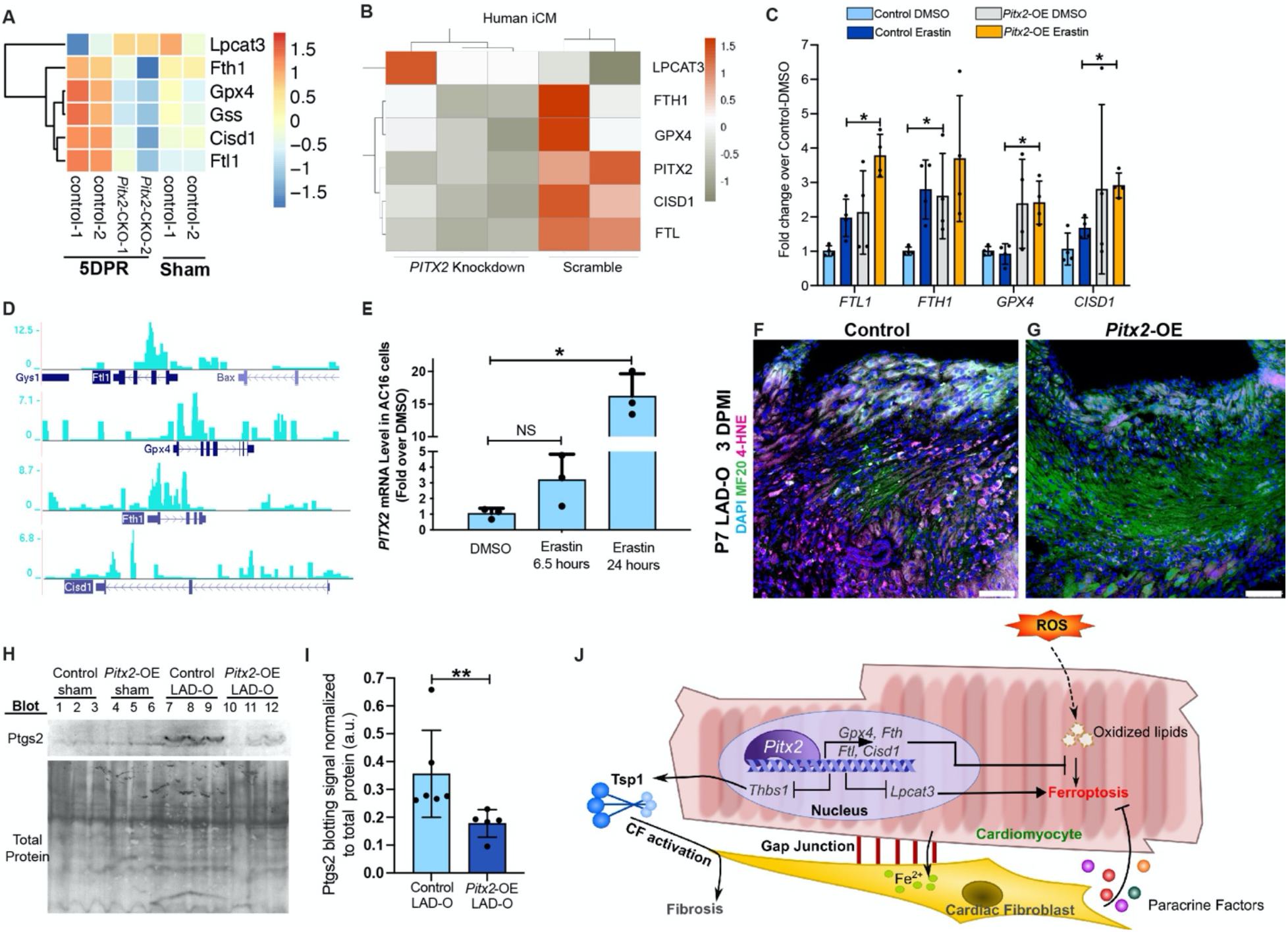
Pitx2 prevents ferroptosis in cardiomyocytes by regulating ferroptotic genes. (A) Heatmap shows transcript level of ferroptotic genes in *Pitx2-CKO (MCK^cre^;Pitx2^f/f^*) ventricles and controls (*Pitx2^f/f^*) at 5 days after apex resection (DPR) or sham. (B) Heatmap shows transcript level of ferroptotic genes in human iCMs after siRNA knockdown of *PITX2*, with scramble controls. (C) qPCR of ferroptotic targets in iCMs after vehicle or erastin treatment, with *PITX2-*expressing or control lentivirus transduction. (D) ChIP-Seq showing Pitx2 binding region near ferroptotic gene loci in regenerating neonatal ventricles. (E) qPCR of *PITX2* in AC16 cells treated with vehicle or erastin. (F, G) Heart tissue of control (F) and *Pitx2-OE* (G) were stained for 4HNE (magenta), MF20 (green) and DAPI (blue) at 3 days after P7 LAD-O. Tamoxifen was administrated daily from P6-8. (H, I) Western blot of Ptgs2 in control and *Pitx2-OE* left ventricles at 3 days after sham or LAD-O at P7. Ptgs2 band intensity quantified in I. (J) Working model of cardiomyocytes interacting with fibroblasts to regulate ferroptosis after MI. Error bars indicate SD. *, p<0.05; **, p<0.01. NS, not significant. Scale bar, 75 μm (F, G).

## DISCUSSION

The discovery of a transient regenerative capacity in neonatal mammalian hearts was an important step in the field of cardiovascular regenerative medicine ^22^, ^43^, ^44^. Neonatal heart regeneration models provide an effective platform for identifying “healing-factors” which could promote adult cardiomyocyte repair ^45^. Here we show that cardiomyocyte survival and the interaction between different cardiac cell types are also important aspects of heart regeneration. After P1 and P7 MI, cardiomyocytes preferably undergo ferroptosis as compared to apoptosis and necroptosis. CFs show resistance to ferroptosis and can protect cardiomyocytes from ferroptosis through paracrine signaling and cell-cell contact. The fibrogenic activity of CFs is kept under control by cardiomyocyte-derived Pitx2 signaling, which also regulates an intrinsic anti-ferroptotic program in cardiomyocytes.

Although cCasp3- or TUNEL-positive cardiomyocytes have been observed in animal models of heart attack, the number of apoptotic cells is minimal ^46^. In a pig model of Ischemia-Reperfusion injury, TUNEL reagent marks roughly 17 cardiomyocytes in every 1 cm^2^ of area ^47^. Therefore, it was imperative to assess alternative RCD in mammalian MI models. Ferroptosis is comparatively new to the family of RCD ^7^. As such, the field has not reached consensus on reliable molecular markers ^41^. The expression of Ptgs2, for example, is associated with ferroptosis, but may not be a part of the driving mechanism ^12^. In fact, the molecular mechanism of ferroptosis may vary based on cell types and diseases, suggesting that cardiomyocytes may have their own specified ferroptotic machinery. Interestingly, our data showed a large quantity of lipid peroxidation accumulated in the nuclear and peri-nuclear area of cardiomyocytes instead of plasma membrane ^48, 49^. This could be a protective mechanism for cardiomyocytes to avoid cell membrane damage and rupture. However, whether cardiomyocytes can efficiently fix the damaged nuclear envelope and other associated membranous structures will require further studies.

CFs, normally considered detrimental in cardiac fibrosis after injury, are required for maintaining the integrity of infarcted myocardium after heart attack ^19^. They promote cardiomyocyte proliferation during development ^50^. The fact that CFs can endure the oxidative microenvironment in an infarcted myocardium makes them candidates as the “guardians” of stressed cardiomyocytes. Here we showed two ways the CFs can protect cardiomyocytes from ferroptosis: the production of protective paracrine factors (including IL-8 and EGF), and the direct interaction that facilitates iron transfer out of cardiomyocytes (Figure 7J).

Interestingly, CFs in both P1 and P7 mouse heart demonstrate resilience to ferroptosis-inducing reagent. They also show high ferritin levels and interact with cardiomyocytes at the border zone of infarcted myocardium. Despite the mouse heart being considered non-regenerative at P7, the protective role of CFs seems to persist. Neonatal mouse hearts can re-establish blood supply to the infarct zone by forming collateral coronary arteries at 4DPMI ^51^. Therefore, ischemic cardiomyocytes, if survive through the initial days after MI, could be rescued and recover. One the other hand, collateral coronary arteries are not formed after P7 LAD-O and more cardiomyocytes will die eventually albeit being protected by CFs.

Importantly, we found that the protective effect of CFs is not restricted to myofibroblasts. Adult cardiomyocytes showed increased level of 4-HNE when resident CFs were removed by genetic approaches. This observation supports the role of fibroblasts in maintaining myocardial homeostasis. The fact that fibroblast-to-myofibroblast transition is not required for this protective mechanism could be taken into consideration when designing anti-fibrotic treatment^52, 53^. Ideally, the anti-ferroptotic function of CFs should not conflict with the anti-fibrotic treatment, which could mainly target the myofibroblasts. The timing of these therapeutic treatment could also be optimized for more favorable outcome.

We previously reported that Pitx2 efficiently promotes myocardial regeneration but had only a mild effect on cardiomyocyte proliferation ^31^. The current studies provided alternative explanations as to how Pitx2 promotes heart repair. In infarcted myocardium, high ROS level and the presence of free iron likely push cardiomyocytes towards a ferroptotic fate. As a key regulator of redox balance, Pitx2 drives an intrinsic anti-ferroptotic mechanism in cardiomyocytes as shown in the current studies. One important hallmark of ferroptotic cells is mitochondrial abnormalities ^7, 8^. We previously reported 505 direct Pitx2 target genes in regenerating neonatal mouse hearts, and Gene Ontology analysis revealed enrichment in mitochondria and respiratory chain genes ^31^. Pitx2 is deeply involved in the maintenance of function and homeostasis of mitochondria, adding more rationale for further dissecting the roles of Pitx2 signaling in ferroptosis. While the CFs protect cardiomyocytes from ferroptosis, their fibrogenic activity is controlled by Pitx2 signaling (Figure 7J). The mechanistic insight into the role of Pitx2 signaling could inform the invention of future therapeutic approaches for treating MI.

## METHODS

### Mouse Lines

All animal protocols and procedures complied with the NIH guidelines and were approved by the Institutional Animal Care and Use Committee (IACUC) of the Medical University of South Carolina (Charleston, South Carolina 29425, USA). The *FVB* (JAX 001800), *C57BL/6J* (JAX 000664), *Mhc^cre-Ert^* (JAX 005657), *Pdgfrα-CreER^T2^* (JAX 032770), *ROSA-DTA* (JAX 009669), *Postn^MCM^* (JAX 029645), ROSA-*mTmG* (JAX 007676) were purchased from the Jackson Laboratory. The *Pitx2^GOF^* transgene for overexpressing *Pitx2* and the floxed allele for *Pitx2 (Pitx2^f/f^*) were described previously ^31, 54^. DNA was extracted from tail or ear biopsies for genotyping using primers provided by JAX or described previously ^31, 54^.

### Left Anterior Descending Coronary Artery Occlusion (LAD-O)

For all mouse survival surgeries, littermate controls were used whenever possible. Both male and female mice were randomly distributed among groups. All surgeries were done blinded from mouse genotype or treatment. LAD-O was performed on P1 and P7 mice as previously described ^24, 31^. Sham procedures were done identically without the occlusion of coronary artery. *Postn^MCM/+^;ROSA-mTmG* mice had LAD-O at P1 or P7; tamoxifen (SigmaAldrich, T5648) was administered subcutaneously at 1, 2, 3 and 5 day-post-surgery before heart tissue collection at 6 days post-surgery. *Postn^MCM/+^;ROSA-DTA* mice were subjected to LAD-O surgery at P1 with tamoxifen administered at P2-4 before heart collection for analysis at P5. *Pitx2-iCKO (Mhc^cre-ERT^;Pitx2^f/f^*) and control (*Mhc^cre-ERT^*) mice were subjected to LAD-O at P1 and tamoxifen injection at P0-2; hearts were collected 3 days post-surgery for analysis. *Pitx2-OE (Mhc^cre-ERT^;Pitx2^GOF^*) and control (*Mhc^cre-ERT^*) mice were subjected to LAD-O at P7 with tamoxifen injection at P6-8, hearts collected at 3 days post-surgery for analysis.

### Echocardiography

8-week-old *Pdgfrα-CreER^T2^;ROSA-DTA* mice had intraperitoneal tamoxifen injection as described in Figure S3H. Echocardiography was performed using a Vevo 3100 ultrasound system (Fujifilm VisualSonics), equipped with a MX550S transducer. B-mode and M-mode data were acquired following the manufacturer’s guidelines.

### Cell culture

HEK293 cells (ATCC CRL-1573) were cultured in Eagle’s Minimum Essential Medium (ATCC, 30-2003), supplemented with 10% FBS (Corning, 35-011-CV) and 1% penicillin/streptomycin (Corning, 30-002-CI). AC16 human cardiomyocyte cell line (MilliporeSigma, SCC109) was cultured in DMEM/Hams F-12 50/50 Mix (Corning, 10-092-CV), supplemented with 12.5%FBS and 1% penicillin/streptomycin. HCF cells (Cell Applications Inc, 306V-05a) were cultured in HCF Growth Medium (Cell Applications Inc, 316-500).

### Differentiation of human iPSCs to cardiomyocytes (iCMs)

iPSCs (Coriell Institute for Medical Research, AICS-0048-039) were cultured in mTeSR1media (STEMCELL Technologies, 85850) on Matrigel (Gibco, A1413302)-coated plates. At 80% of confluency, iPSCs were differentiated into iCMs as previously described ^26^. Briefly, the iPSCs were treated with 8μM CHIR-99021 (SelleckChem, S2924) in RPMI (Gibco, 11875093)-B27(no insulin) (Gibco, A1895601) from day 0-1. Media was changed on day 2 and the cells were treated with 5 μM IWR1 (SelleckChem, S7086) in RPMI-B27(no insulin) from day 3-4. Starting from day 7, RPMI-B27(with insulin) (Gibco, 17504044) media were given to iPSC-derived cardiomyocytes (iCMs). Two rounds of glucose starvation from day 12-15 and from day 20-23 were performed to eliminate non-cardiomyocyte cells. At day 30, iCMs were transferred to Matrigel-coated 24-well plates and maintained for further use. For co-culture of iCM and HCF, iCM cells were seeded at 2.5×10^4^ cells per well in a Matrigel-coated 24-well plate on day 1, and HCF cells were seeded at 1.5×10^4^ cells per well on day 2. Cells were cultured in RPMI-B27 (with insulin). Co-culture was used on day 3 for experiments and analysis.

### Primary culture of mouse cardiac fibroblast

Isolated wild type hearts (P1 or P7) were minced into 1 mm^3^ pieces and placed in 0.1% gelatin (Sigma, G1890)-coated 6-well plates with 300 μl explant media (DMEM, 20%FBS). Three hours later, when tissue pieces settled down, another 2 ml media was slowly added and was replaced every three days. Migrated cells were harvested at day 7 and filtered through 40 μm cell strainers for further experiments ^32^.

### HCF-conditioned medium

HCF cells were cultured in 10-cm dishes to confluency, washed with PBS, and then treated with 200 μM H_2_O_2_ in DMEM basal media. After 2 hours, cells were washed with PBS for three times and cultured in DMEM basal media overnight. Conditioned media were collected and filtered through a 0.2 μm Supor^®^ membrane (Pall Laboratory, 4612). Control conditioned medium was prepared in the same way but treated with H_2_O instead of H_2_O_2_.

### Cytokine array assay

HCF-conditioned media were applied to the Proteome Profiler Human XL Cytokine Array Kit (R&D Systems, ARY022B). 5ml of conditioned medium from H_2_O_2_ or H_2_O treated HCFs was concentrated into 500μl using MilliporeSigma Protein-Concentrate Kit (Fisher Scientific, 50-525-36) according to the manufacturer’s instructions. Equal amount of total protein from all groups was loaded on the cytokine array. Blotting signal was visualized using chemiluminescence detection and a ChemiDoc Touch Imaging System (Bio-Rad, 12003154).

### Living imaging of HCF and iCM co-culture with iron overload

iCMs were seeded and allowed to acclimate for 3 days until spontaneously beating was observed. siRNA was used to knockdown *CX43* and *CX45* for 24 hours. iCMs were then washed with PBS. Similar number of HCFs were added to iCM culture to reach confluency. The co-culture was allowed to acclimate overnight. The following morning cells were washed with PBS and treated with FluoroBrite DMEM media containing 25 μM of FAS (ferrous ammonium sulfate) for 1 hour. Following treatment cells were washed with PBS for three times and BioTracker FerroOrange Live Cell Dye (MilliporeSigma, SCT210) (1000×) diluted in FluoroBrite DMEM was added to the cells. After 1 hour of incubation, cells were washed twice with FluoroBrite DMEM and imaged on a Leica SP8 confocal microscope with live imaging chamber.

### Transfection of siRNA in HCF and iCM cells

Lipofectamine RNAiMAX transfection reagent (ThermoFisher Scientific, 13778030) was used to deliver siRNA into HCFs and iCMs following the manufacturer’s guidelines. Scrambled siRNA was used as control. The siRNA oligonucleotides were pre-designed DsiRNA Duplex from Integrated DNA Technologies (IDT) with oligonucleotide sequences as follow:

*FTH1* (hs.Ri.FTH1.13.2)
rGrUrUrUrArCrCrUrGrUrCrCrArUrGrUrCrUrUrArCrUrAC T rArGrUrArGrUrArArGrArCrArUrGrGrArCrArGrGrUrArArArCrGrU
*FTL* (hs.Ri.FTL.13.1)
rGrArGrGrArArGrUrGrArArGrCrUrUrArUrCrArArGrArAGA rUrCrUrUrCrUrUrGrArUrArArGrCrUrUrCrArCrUrUrCrCrUrCrArU
*CX43* (hs.Ri.GJC1.13.1) rArGrArArUrGrGrArCrUrUrArCrArUrGrUrUrArUrCrArGT T rArArCrUrGrArUrArArCrArUrGrUrArArGrUrCrCrArUrUrCrUrUrC
*CX45* (hs.Ri.GJA1.13.2)
rGrUrGrGrUrArCrArUrCrUrArUrGrGrArUrUrCrArGrCrUTG rCrArArGrCrUrGrArArUrCrCrArUrArGrArUrGrUrArCrCrArCrUrG Scrambled negative control rCrUrUrCrCrUrCrUrCrUrUrUrCrUrCrUrCrCrCrUrUrGrUGA rUrCrArCrArArGrGrGrArGrArGrArArArGrArGrArGrGrArArGrGrA

### Lentivirus transduction

Coding sequence of human *PITX2C* (NM_000325.6) was cloned into lentivirus transfer plasmid (pWPI, Addgene#12254) using Gibson Assembly^®^ kit (NEB, E2611). Primers used for amplifying *PITX2C* insert Forward: 5’-CTAGCCTCGAGGTTTAT GAACTGCATGAAAGGCCC-3’, reverse: 5’-TGCAGCCCGTAGTTTCCGGCAAGG TCCTAGGATCC-3’. Empty pWPI was used for preparing control lentivirus. For lentivirus packaging, pWPI, pMD2.G (Addgene#12259) and psPAX2 (Addgene#12260) plasmids were co-transfected into Lenti-X^TM^ 293T cells (Clontech Laboratories, 632180) using Lipofectamine 3000 (Fisher, L3000008). Virus particles in the supernatant was collected, filtered and stored in −80°C freezer until be used.

### Cell viability measurement

For low density, HCF, HEK293, and iCM cells were seeded at 3×10^3^ cells per well in a 24-well plate. For middle density, HCF and HEK293 cells were seeded at 3×10^4^ cells per well while iCM cells were seeded at 2×10^4^ per well in a 24-well plate. For high density, HCF and HEK293 cells were seeded at 10^5^ cells per well and iCMs were seeded at 6×10^4^ cells per well in 24-well plate (Figure S3A). Cells were cultured overnight and then treated with erastin (Sigma, E7781) at varying concentrations in DMEM or RPMI basal medium for 5 or 16 hours. DMSO-treated wells were used as controls. HCF and HEK293 cells were treated with varying concentrations of H_2_O_2_ for 6 hours in DMEM basal medium and iCMs were treated similarly but in RPMI basal medium. Primary culture of P1 and P7 mouse cardiac fibroblasts were treated with erastin or DMSO for 16 hours in DMEM basal medium. Cell viability was assessed using Trypan Blue (Invitrogen, T10282) according to the manufacturer’s instructions. Cell viability was reported as a ratio of Trypan-Blue-negative cell number in experimental group over the vehicle control group. AC16 cells were seeded at low (3×10^4^) or high (8×10^4^) density in a 24-well plate and cultured overnight before being treated with erastin at varying concentrations in F12/DMEM basal medium, with DMSO as controls for 9 hours. Cells were then washed with HEPES then stained with DAPI and Sytox Green (Thermo Fisher, S7020) at 37°C for 20 minutes. Following staining, cells were washed with HEPES and immediately imaged using a Lionheart FX Automated Live Cell Imager (Agilent).

### Fe^2+^ staining

Cells were stained with FerroOrange Live Cell Dye according to the manufacturer’s guideline. Briefly, cells were rinsed with PBS. The dye was pre-diluted and added to the cells. After 30 minutes of incubation at 37°C, cells were rinsed with PBS twice and fixed in 10% Neutral Buffered Formalin (Leica, 3800598) for 15 min at room temperature before imaging using a Leica SP8 confocal microscope.

### Tissue processing, histology, and immunohistochemistry

Hearts were fixed in 10% formalin (VWR, 10015-192) at room temperature overnight with continuous rocking. The tissue was then processed for paraffin embedding. Tissue was sectioned at 7 μm thickness, deparaffinized in xylene, rehydrated and subjected to histology or immunofluorescence staining. Tissue slides were incubated in humid chamber overnight at 4°C in primary antibody in 1% BSA in PBS. On the second morning, slides were washed in PBS then incubated for 1 hour at room temperature in secondary antibody in 1% BSA in PBS. Slides were washed with PBS and counterstained with DAPI (Sigma, D9542) then mounted in VECTASHIELD hardset mounting medium (Vector Laboratories, H1400). HCFs were fixed in formalin for 15 minutes followed by PBS wash and permeabilization in 0.2% Triton X-100 (Sigma, T8787) in PBS. Cells were blocked for 1 hour in 5% BSA (Sigma, A3059) in PBS at room temperature before the incubation with primary antibody for 2 hours and fluorophore-labeled secondary antibody for 1 hour. Cells were counterstained with DAPI (Sigma, D9542) and placed in cold PBS for imaging. Immunofluorescent images were acquired using a Leica SP8 confocal microscope (Leica Microsystems). Primary antibodies used include: cleaved Caspase 3 (1:200, Cell Signaling, 9664), pMLKL (1:100, Cell Signaling, D6E3G), PTGS2 (1:100, Proteintech, 12375-1-AP), 4-HNE (1:100, Bioss, bs-6313R), αSMA (1:400, Sigma, C6198), pSMAD2 (1:200, Abcam, ab188334), pSMAD3 (1:200, Abcam, ab52903), TGFβRII (1:200 Abcam, ab186838), FTH1 (1:200, Abcam, ab183781), FTL (1:100, Abcam, ab109373) CX45 (1:200, Abcam, ab135474), CX43 (1:200, Sigma, C6219), PDGFRα (1:100, R&D Systems, AF1062-SP), MF20 (1:400, Developmental Studies Hybridoma Bank, AB_2147781), Vimentin (1:400, Abcam, ab92547), TSP1 (1:100, Abcam, ab85762), and cTNT (1:400, Fisher, ms-295p1abx). Secondary antibodies used include: Goat Anti-Mouse IgG 488 (1:400, Santa Cruz Biotechnology, A32723), Chicken Anti-Mouse IgG 488 (1:400, Thermo Invitrogen, A21200), Donkey Anti-Rabbit IgG 647 (1:400, Thermo Invitrogen, A31573), Alexa 488 Goat Anti-Rabbit (1:400, Thermo Invitrogen, A11008), Donkey Anti-Goat IgG 633 (1:400, Thermo Invitrogen, A21082), Alexa 633 Goat Anti-Mouse (1:400, Thermo Invitrogen, A21050), Donkey Anti-Goat 568 (1:200, Thermo Invitrogen, A11057). For Picro-Sirius Red staining, tissue slides were deparaffinized, washed with water, before immersion in 0.1% Picro-Sirius Red made with picric acid (Sigma, P6744-1GA) and Direct Red (Sigma, 365548). Slides were then washed in 0.5% glacial acetic acid (Sigma, A6283), dehydrated in ethanol, cleared in xylene, and mounted using Shurmount (General Data, LC-A).

### Western blot

HCFs or mouse heart tissue were lysed in RIPA buffer. Protein concentration was determined using Pierce BCA protein assay kit (Thermo Fisher, PI23227) according to manufacturer’s instructions. Western blot was performed as previously described ^55^. Total protein was examined using Pierce Reversible Protein Stain Kit for PVDF membranes (Thermo Fisher, PI24585) and imaged using Bio-Rad ChemiDoc Imaging System. Target protein was detected using SuperSignal West Pico Chemiluminescent Substrate (Thermo Fisher, 34577). Target band intensities were quantified using ImageJ software (National Institutes of Health). Primary Antibodies used include: FTH1 (1:1000, Santa Cruz Biotechnology, sc-376594), FTL (1:1000, Abcam, ab109373), PTGS2 (1:1000, Proteintech, 12375-1-AP), TSP1 (1:1000, Abcam, ab85762), and α-TUBULIN (1:1000, Sigma, T5168-100UL). Secondary antibodies include goat-anti-mouse (1:2500, Thermo Invitrogen, 32430) or goat-anti-rabbit (1:2500, Santa Cruz Biotechnology, sc-2357) horseradish peroxidase (HRP)-conjugated antibodies.

### qPCR

RNA was prepared from cells or heart tissue using TRIzol™ (Fisher, 15596018) according to manufacturer’s guideline. cDNA was prepared from 1 μg of RNA with iScript Reverse Transcription Supermix (Bio-Rad, 1708841). qPCR was performed using SsoAdvanced Universal SYBR Green Supermix (Bio-Rad, 172-5271) and a CFX96 Touch Real-Time PCR machine (Bio-Rad). Primers used were as follows: *PITX2*, forward: 5’-AGCGGACTCACTTTACCAGC-3’, reverse: 5’-CCGTAAGGTTGGTCCACACA-3’; *FTL*, forward: 5’-TGGGCTTCTATTTCGACCGC-3’, reverse: 5’-TTTCATGGCGTCTGGGGTTT-3’; *FTH1*, forward: 5’-GCTCTACGCCTCCTACGTTT-3’, reverse: 5’-AAGGAAGATTCGGCCACCTC-3’; *CISD1*, forward: 5’-GCTGTGTACTGCCGTTGTTG-3’, reverse: 5’-TGATCAGAGGGCCCACATTG-3’; *GPX4*, forward: 5’-GAGGCAAGACCGAAGTAAACTAC-3’, reverse: 5’-CCGAACTGGTTACACGGGAA-3’; *ACTB*, forward: 5’-CAATGAGCTGCGTGTGGCT-3’, reverse: 5’-GGATAGCACAGCCTGGATAGCAA-3’; *HSP90*, forward: 5’-CGAAGTTGGACAGTGGTAAAGAG-3’, reverse: 5’-TGCCCAATCATGGAGATGTCT-3’; *Cxcr1*, forward: 5’-TGTCCCTTCTGAGCTTGCTG-3’, reverse: 5’-CCAAGAAGGGCAGGGTCAAT-3’; *Cxcr2*, forward: 5’-CGCTGCTCATCATGCTGTTC-3’, reverse: 5’-GCAGGAAGACAAGGACGACA-3’.

### RNA-Seq

Total RNA was extracted from iCMs treated with siRNA targeting *PITX2* or scramble control using TRIzol™ Reagent. RNA-Seq was performed using Novogene’s Illumina NovaSeq PE150 for nextgeneration sequencing according to the manufacturer’s direction. Sequenced reads in Fastq format were aligned to *mm10* reference genome using STAR (Spliced Transcripts Alignment to a Reference) alignment software ^56^. R package DESeq2 was used to normalize and quantify the aligned RNA-seq reads with threshold P ≤ 0.05 and log2foldchange ≤ −0.58 to compare each treatment groups to another ^57^. Heatmaps were created using R package Pheatmap. The RNA-Seq data from this study have been deposited in GEO under accession number GSE211568.

### Statistics

All quantitative experiments (qPCR, western blot, cell count) included at least 3 biological replicates. Animal studies included at least 3 mice per group. Statistical significance was determined by t-test. Equal variance was assumed. Outliers were determined by Grubb’s test. All statistical work was done on GraphPad Prism (GraphPad Software). All bar graphs included scattered dots. All bar graphs represent mean ± SD. *P<0.05, **P<0.01, ***P<0.001, and ****P<0.0001 were considered statistically significant.

## SUPPLEMENTAL INFORMATION

There are 6 Supplemental figures.

## ACKNOWLEDGMENTS

This project was supported by grants from the National Institutes of Health (NIH) (1R01HL148728 to G.T.; HL144938 to H.M.S). G.T. was supported in part by American Heart Association (AHA) (17SDG33400141), Saving Tiny Hearts Society, NSF EPSCoR RII Track-1: Materials Assembly and Design Excellence in South Carolina (MADE in SC) OIA-1655740, and NIH (1R21AI162775-01A1 / 412712-19270). M.E.M. was supported in part by training grant from NIH (R25GM072643). A.M.T. was supported in part by training grant from NIH (T32GM132055). R.A.S was supported in part by training grant from NIH (HL007260). C.L. was supported by AHA (19CDA34760019). M.Z. was supported by National Natural Science Foundation of China (81901489). The Molecular Analytics Core at the Medical University of South Carolina (MUSC) is supported by NIGMS GM103499 and MUSC’s Office of the Vice President for Research. We thank all members of the Tao and Sucov laboratories for their constructive feedback during the preparation of manuscript. We thank the husbandry and veterinary staff at MUSC Division of Laboratory Animal Resources. We also thank the Department of Regenerative Medicine and Cell Biology at the MUSC for providing common equipment.

## AUTHOR CONTRIBUTIONS

G.T. conceived and supervised the project. G.T., M.E.M., S.L. designed the experiments. M.E.M, S.L., G.T., A.M.T., R.A.S., P.L.R. and C.L. performed experiments and analyzed data. H.M.S. provided reagents and scientific advice. C.L. provided reagents and advice on hiPSC platform. A.M.T., G.T. and M.Z. performed bioinformatics and statistical analyses. G.T., M.E.M., S.L. and A.M.T. wrote the manuscript.

## REFERENCES

1. Benjamin, E.J., et al., Heart Disease and Stroke Statistics-2019 Update: A Report From the American Heart Association. Circulation, 2019. 139(10): p. e56–e528.

2. Lopaschuk, G.D., R.L. Collins-Nakai, and T. Itoi, Developmental changes in energy substrate use by the heart. Cardiovasc Res, 1992. 26(12): p. 1172–80.

3. Puente, B.N., et al., The oxygen-rich postnatal environment induces cardiomyocyte cellcycle arrest through DNA damage response. Cell, 2014. 157(3): p. 565–79.

4. Del Re, D.P., et al., Fundamental Mechanisms of Regulated Cell Death and Implications for Heart Disease. Physiol Rev, 2019. 99(4): p. 1765–1817.

5. Liu, K., et al., A double-edged sword: role of apoptosis repressor with caspase recruitment domain (ARC) in tumorigenesis and ischaemia/reperfusion (I/R) injury. Apoptosis, 2023.

6. Maslov, L.N., et al., The regulation of necroptosis and perspectives for the development of new drugs preventing ischemic/reperfusion of cardiac injury. Apoptosis, 2022. 27(9-10): p. 697–719.

7. Dixon, S.J., et al., Ferroptosis: an iron-dependent form of nonapoptotic cell death. Cell, 2012. 149(5): p. 1060–72.

8. Friedmann Angeli, J.P., et al., Inactivation of the ferroptosis regulator Gpx4 triggers acute renal failure in mice. Nat Cell Biol, 2014. 16(12): p. 1180–91.

9. Stockwell, B.R., et al., Ferroptosis: A Regulated Cell Death Nexus Linking Metabolism, Redox Biology, and Disease. Cell, 2017. 171(2): p. 273–285.

10. Fang, X., et al., Loss of Cardiac Ferritin H Facilitates Cardiomyopathy via Slc7a11-Mediated Ferroptosis. Circ Res, 2020. 127(4): p. 486–501.

11. Bulluck, H., et al., Residual Myocardial Iron Following Intramyocardial Hemorrhage During the Convalescent Phase of Reperfused ST-Segment-Elevation Myocardial Infarction and Adverse Left Ventricular Remodeling. Circ Cardiovasc Imaging, 2016. 9(10): p. e004940.

12. Yang, W.S., et al., Regulation of ferroptotic cancer cell death by GPX4. Cell, 2014. 156(1-2): p. 317–331.

13. Bersuker, K., et al., The CoQ oxidoreductase FSP1 acts parallel to GPX4 to inhibit ferroptosis. Nature, 2019. 575(7784): p. 688–692.

14. Doll, S., et al., FSP1 is a glutathione-independent ferroptosis suppressor. Nature, 2019. 575(7784): p. 693–698.

15. Fang, X., et al., Ferroptosis as a target for protection against cardiomyopathy. Proc Natl Acad Sci U S A, 2019. 116(7): p. 2672–2680.

16. Baba, Y., et al., Protective effects of the mechanistic target of rapamycin against excess iron and ferroptosis in cardiomyocytes. Am J Physiol Heart Circ Physiol, 2018. 314(3): p. H659–H668.

17. Wu, J., et al., Intercellular interaction dictates cancer cell ferroptosis via NF2-YAP signalling. Nature, 2019. 572(7769): p. 402–406.

18. Fu, X., et al., Specialized fibroblast differentiated states underlie scar formation in the infarcted mouse heart. J Clin Invest, 2018. 128(5): p. 2127–2143.

19. Kanisicak, O., et al., Genetic lineage tracing defines myofibroblast origin and function in the injured heart. Nat Commun, 2016. 7: p. 12260.

20. Vasquez, C., et al., Enhanced fibroblast-myocyte interactions in response to cardiac injury. Circ Res, 2010. 107(8): p. 1011–20.

21. Miragoli, M., G. Gaudesius, and S. Rohr, Electrotonic modulation of cardiac impulse conduction by myofibroblasts. Circ Res, 2006. 98(6): p. 801–10.

22. Porrello, E.R., et al., Transient regenerative potential of the neonatal mouse heart. Science, 2011. 331(6020): p. 1078–80.

23. Bergmann, O., et al., Dynamics of Cell Generation and Turnover in the Human Heart. Cell, 2015. 161(7): p. 1566–75.

24. Heallen, T., et al., Hippo signaling impedes adult heart regeneration. Development, 2013. 140(23): p. 4683–90.

25. Sun, L., et al., Mixed lineage kinase domain-like protein mediates necrosis signaling downstream of RIP3 kinase. Cell, 2012. 148(1-2): p. 213–27.

26. Lian, X., et al., Directed cardiomyocyte differentiation from human pluripotent stem cells by modulating Wnt/beta-catenin signaling under fully defined conditions. Nat Protoc, 2013. 8(1): p. 162–75.

27. Dolma, S., et al., Identification of genotype-selective antitumor agents using synthetic lethal chemical screening in engineered human tumor cells. Cancer Cell, 2003. 3(3): p. 285–96.

28. Davidson, M.M., et al., Novel cell lines derived from adult human ventricular cardiomyocytes. J Mol Cell Cardiol, 2005. 39(1): p. 133–47.

29. Davis, J., et al., A TRPC6-dependent pathway for myofibroblast transdifferentiation and wound healing in vivo. Dev Cell, 2012. 23(4): p. 705–15.

30. Chung, M.I., et al., Niche-mediated BMP/SMAD signaling regulates lung alveolar stem cell proliferation and differentiation. Development, 2018. 145(9).

31. Tao, G., et al., Pitx2 promotes heart repair by activating the antioxidant response after cardiac injury. Nature, 2016. 534(7605): p. 119–23.

32. Wang, L., et al., Improved Generation of Induced Cardiomyocytes Using a Polycistronic Construct Expressing Optimal Ratio of Gata4, Mef2c and Tbx5. J Vis Exp, 2015 (105).

33. Harada, A., et al., Essential involvement of interleukin-8 (IL-8) in acute inflammation. J Leukoc Biol, 1994. 56(5): p. 559–64.

34. Bersell, K., et al., Neuregulin1/ErbB4 signaling induces cardiomyocyte proliferation and repair of heart injury. Cell, 2009. 138(2): p. 257–70.

35. Holmes, W.E., et al., Structure and functional expression of a human interleukin-8 receptor. Science, 1991. 253(5025): p. 1278–80.

36. Suarez-Ortegon, M.F., et al., Ferritin, metabolic syndrome and its components: A systematic review and meta-analysis. Atherosclerosis, 2018. 275: p. 97–106.

37. Chen, X., et al., Iron Metabolism in Ferroptosis. Front Cell Dev Biol, 2020. 8: p. 590226.

38. Brown, T.R., T. Krogh-Madsen, and D.J. Christini, Illuminating Myocyte-Fibroblast Homotypic and Heterotypic Gap Junction Dynamics Using Dynamic Clamp. Biophys J, 2016. 111(4): p. 785–797.

39. Harris, A.L., Connexin channel permeability to cytoplasmic molecules. Prog Biophys Mol Biol, 2007. 94(1-2): p. 120–43.

40. Crawford, S.E., et al., Thrombospondin-1 is a major activator of TGF-beta1 in vivo. Cell, 1998. 93(7): p. 1159–70.

41. Tang, D., et al., Ferroptosis: molecular mechanisms and health implications. Cell Res, 2021. 31(2): p. 107–125.

42. Dixon, S.J., et al., Human Haploid Cell Genetics Reveals Roles for Lipid Metabolism Genes in Nonapoptotic Cell Death. ACS Chem Biol, 2015. 10(7): p. 1604–9.

43. Ye, L., et al., Early Regenerative Capacity in the Porcine Heart. Circulation, 2018. 138(24): p. 2798–2808.

44. Zhu, W., et al., Regenerative Potential of Neonatal Porcine Hearts. Circulation, 2018. 138(24): p. 2809–2816.

45. Deshmukh, V., J. Wang, and J.F. Martin, Leading progress in heart regeneration and repair. Curr Opin Cell Biol, 2019. 61: p. 79–85.

46. van Empel, V.P., et al., Myocyte apoptosis in heart failure. Cardiovasc Res, 2005. 67(1): p. 21–9.

47. Weil, B.R., et al., Brief Myocardial Ischemia Produces Cardiac Troponin I Release and Focal Myocyte Apoptosis in the Absence of Pathological Infarction in Swine. JACC Basic Transl Sci, 2017. 2(2): p. 105–114.

48. Doll, S., et al., ACSL4 dictates ferroptosis sensitivity by shaping cellular lipid composition. Nat Chem Biol, 2017. 13(1): p. 91–98.

49. Riegman, M., et al., Ferroptosis occurs through an osmotic mechanism and propagates independently of cell rupture. Nat Cell Biol, 2020. 22(9): p. 1042–1048.

50. Ieda, M., et al., Cardiac fibroblasts regulate myocardial proliferation through beta1 integrin signaling. Dev Cell, 2009. 16(2): p. 233–44.

51. Das, S., et al., A Unique Collateral Artery Development Program Promotes Neonatal Heart Regeneration. Cell, 2019. 176(5): p. 1128–1142 e18.

52. Qian, L., et al., In vivo reprogramming of murine cardiac fibroblasts into induced cardiomyocytes. Nature, 2012. 485(7400): p. 593–8.

53. Aghajanian, H., et al., Targeting cardiac fibrosis with engineered T cells. Nature, 2019. 573(7774): p. 430–433.

54. Wang, J., et al., Pitx2 prevents susceptibility to atrial arrhythmias by inhibiting left-sided pacemaker specification. Proc Natl Acad Sci U S A, 2010. 107(21): p. 9753–8.

55. Li, L., et al., Pitx2 maintains mitochondrial function during regeneration to prevent myocardial fat deposition. Development, 2018.

56. Dobin, A., et al., STAR: ultrafast universal RNA-seq aligner. Bioinformatics, 2013. 29(1): p. 15–21.

57. Love, M.I., W. Huber, and S. Anders, Moderated estimation of fold change and dispersion for RNA-seq data with DESeq2. Genome Biol, 2014. 15(12): p. 550.

