## Supplemental figures and legend for "Cardiomyocyte-fibroblast interaction regulates ferroptosis and fibrosis after myocardial injury"

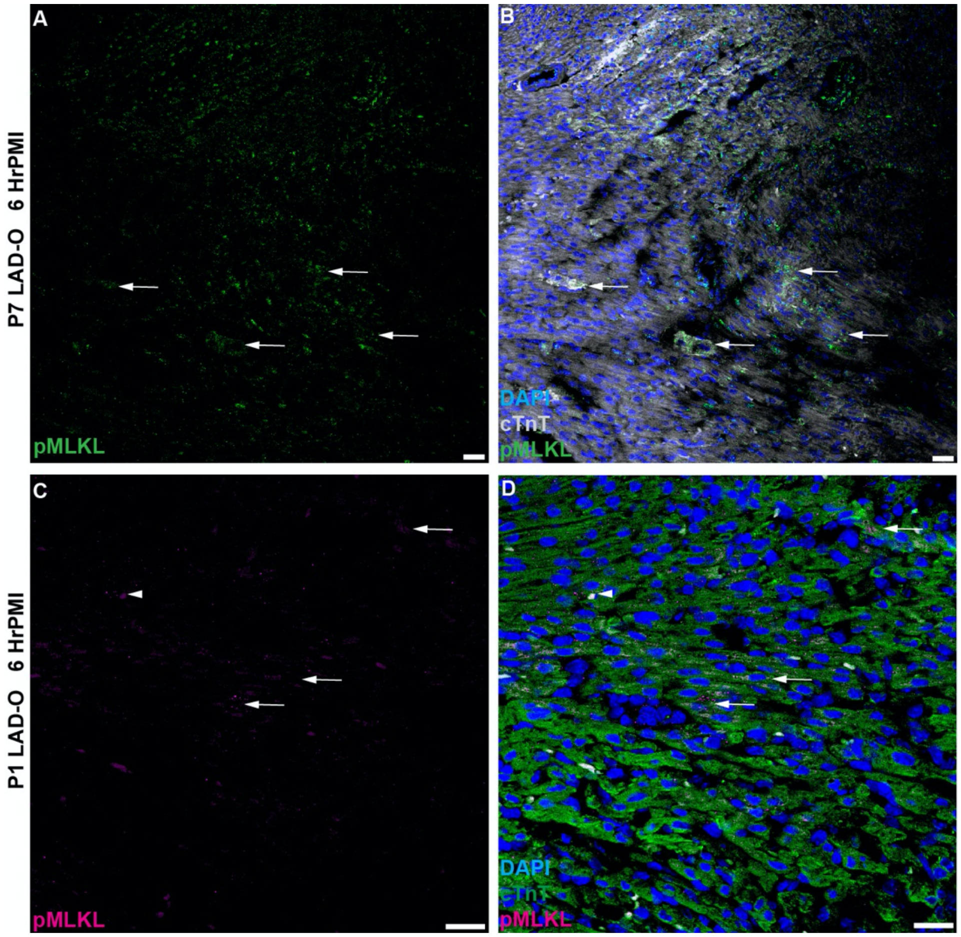


**Figure S1. Necroptosis in cardiomyocytes immediately after MI, related to Figure 1.** (A, B) Heart tissue section of wild type mouse stained for pMLKL (Green) and cTnT (grey) at 6 hours after P7 LAD-O. (C, D) Heart tissue section of wild type mouse stained for pMLKL (magenta) and cTnT (green) at 6 hours after P1 LAD-O. DAPI in blue. Arrows, cardiomyocytes positive for pMLKL. Scale bar, 25 μm.

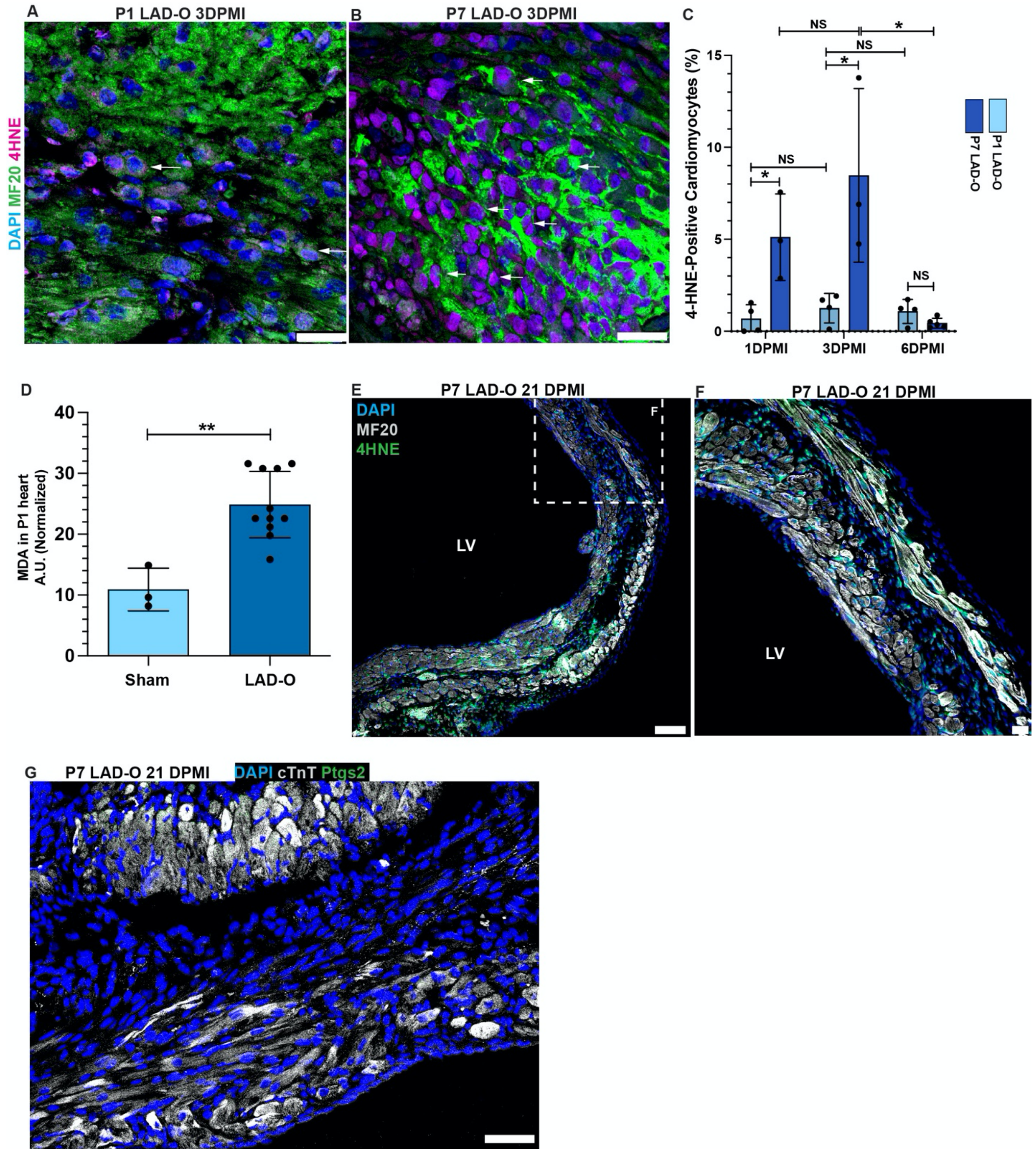


**Figure S2. Ferroptosis occurs in cardiomyocytes after MI, related to Figure 1.** (A, B) Mouse heart tissue stained for 4-HNE (magenta) and MF20 (green) at 3 days after P1 (A) or P7 (B) LAD-O. Arrows, cardiomyocytes positive for 4-HNE. (C) Ratio of cardiomyocytes positive for 4-HNE at 1, 3 and 6 days after P1 or P7 LAD-O. (D) MDA assay quantified lipid peroxidation level in ventricular myocardium after P1 LAD-O or sham procedure. (E, F) Mouse heart tissue stained for 4-HNE (green) and MF20 (grey) at 21 days after P7 LAD-O. (G) Heart tissue stained for Ptgs2 (green) and MF20 (grey) at 21 days after P7 LAD-O. LV, left ventricle. DAPI in blue. Error bars indicate SD. *, $p<0.05$. **, $p<0.01$. NS, not significant. Scale bar, 25 $\mu$m (A, B, F, G), 100 $\mu$m (E).

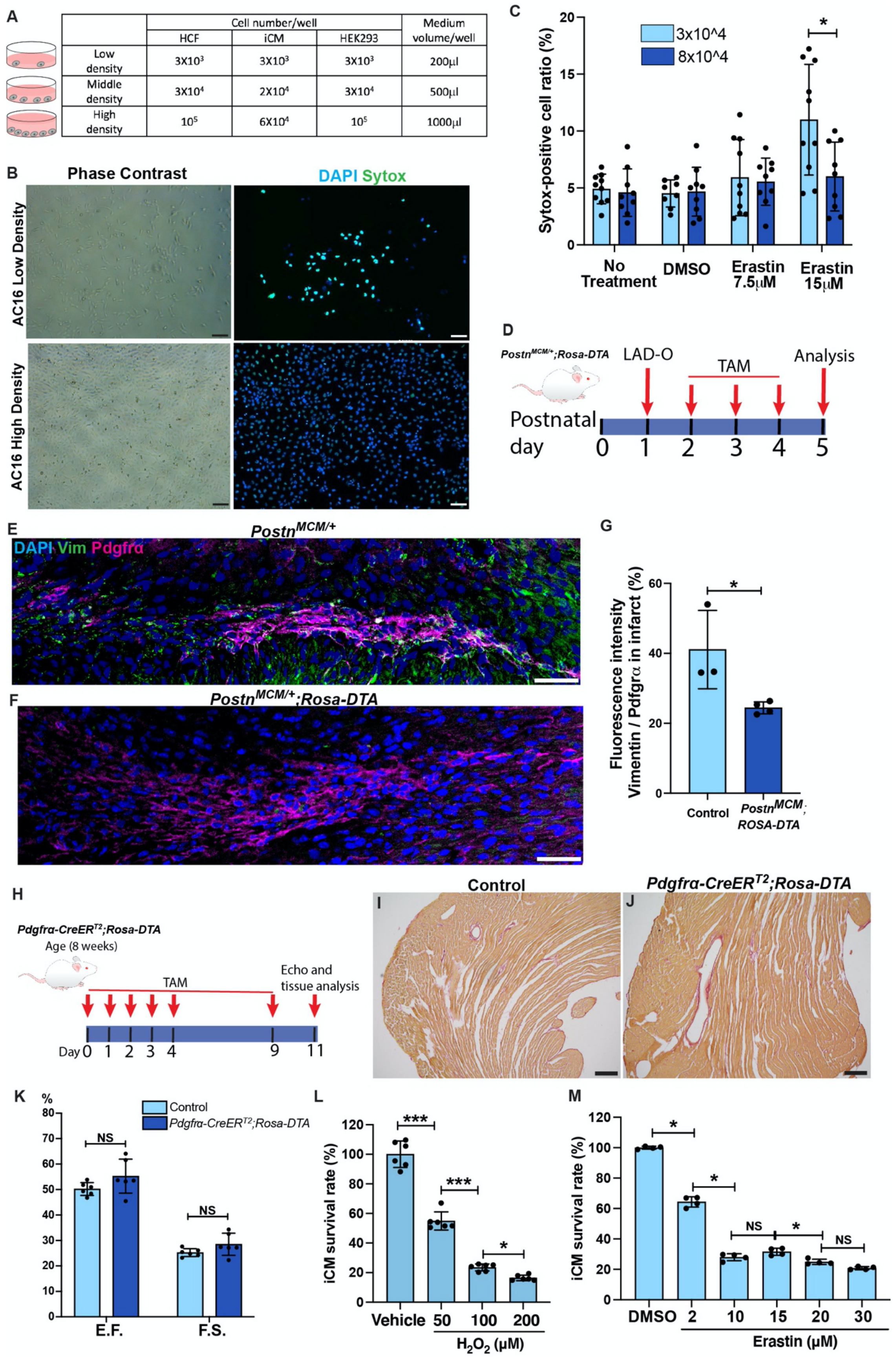
A
Cell number/well
HCF
iCM
HEK293
Medium volume/well
Low density
3X10³ 3X10³ 3X10³ 200µl
Middle density
3X10⁴ 2X10⁴ 3X10⁴ 500µl
High density
10⁵ 6X10⁴ 10⁵ 1000µl
B
Phase Contrast
DAPI Sytox
AC16 Low Density
AC16 High Density
C
3x10^4
8x10^4
Sytox-positive cell ratio (%)
No Treatment
DMSO
Erastin 7.5µM
Erastin 15µM
D
Postn^MCM/+;Rosa-DTA
LAD-O
TAM
Analysis
Postnatal day
0 1 2 3 4 5
E
Postn^MCM/+
DAPI Vim Pdgfrα
F
Postn^MCM/+;Rosa-DTA
G
Fluorescence intensity Vimentin / Pdgfrα in infarct (%)
Control
Postn^MCM;ROSA-DTA
H
Pdgfrα-CreER^T2;Rosa-DTA
Age (8 weeks)
TAM
Echo and tissue analysis
Day 0 1 2 3 4 9 11
Control
Pdgfrα-CreER^T2;Rosa-DTA
I
J
K
%
Control
Pdgfrα-CreER^T2;Rosa-DTA
NS
E.F.
F.S.
L
iCM survival rate (%)
Vehicle 50 100 200
H₂O₂ (µM)
M
iCM survival rate (%)
DMSO 2 10 15 20 30
Erastin (µM)
NS

**Figure S3. Cell density regulates ferroptosis *in vitro* and *in vivo*, related to Figure 2 and Figure 3.**

(A) Schematic of cell density assays, see Figure 2 and 3. (B) AC16 cells cultured at low ($3\times10^4$/well) and high ($8\times10^4$/well) density and treated with erastin. Dying cells were stained with Sytox (green). (C) Ratio of Sytox-positive AC16 cells. (D) Schematic plan for E-G and Figure 2H-P. (E, F) Heart tissue of controls (*Postn*$^{MCM/+}$, E) and *Postn*$^{MCM/+}$*;ROSA-DTA* (F) mice were stained for Vim (green) and Pdgfr$\alpha$ (magenta) at 4 DPMI after P1 LAD-O. (G) Ratio of fluorescence intensity, Vim over Pdgfr$\alpha$. (H) Schematic plan for I-K and Figure 2R-W. (I, J) Picrosirius red staining of adult control (*Rosa-DTA*, I) and *Pdgfr$\alpha$-CreER*$^{T2}$*;ROSA-DTA* (J) mice. (K) Ejection fraction (E.F.) and fractional shorting (F.S.) of control and *Pdgfr$\alpha$-CreER*$^{T2}$*;ROSA-DTA* mice. (L) Survival rate of iCM after treatment with 50, 100 or 200 μM of $H_2O_2$, compared to vehicle ($H_2O$). (M) Survival rate of iCMs after erastin treatment at 2, 10, 15, 20 or 30 μM, compared to DMSO control. Nuclei stained with DAPI (blue). TAM, tamoxifen. Error bars indicate SD. *, $p<0.05$. ***, $p<0.001$. NS, not significant. Scale bar, 100 μm (B, I, J), 25 μm (E, F).

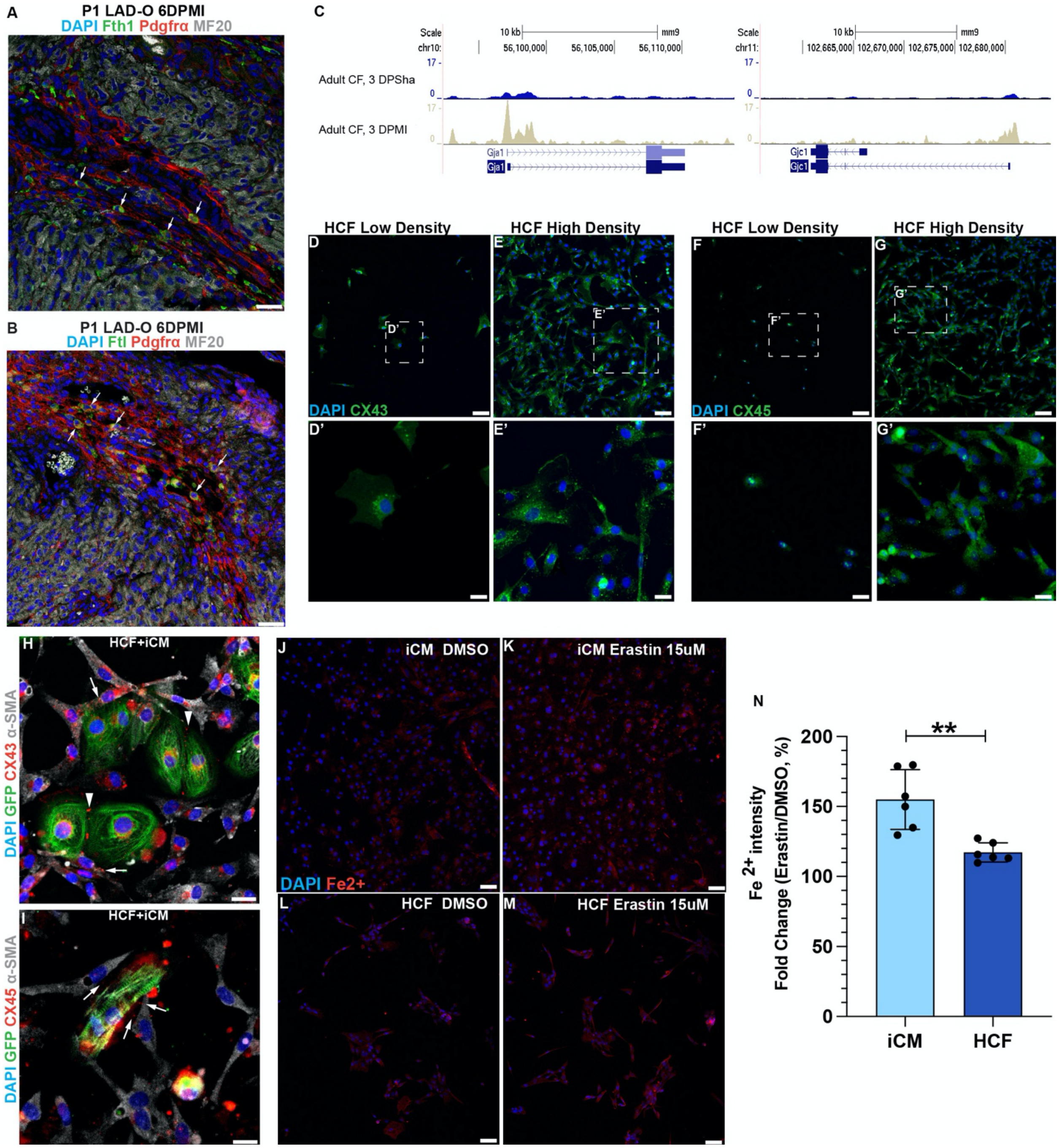


**Figure S4. Cardiac fibroblasts interact with cardiomyocytes to share free iron, related to Figure 5.** (A, B) Mouse heart sections stained for Fth1 (green, A) or Ftl (green, B), with Pdgfrα (red) and MF20 (grey) at 6 days after P1 LAD-O. Arrows, Pdgfrα-labeled cells positive for Fth1 (A) or Ftl (B). (C) ATAC-Seq shows open chromatin region at loci of *Gja1* (*Cx4*3) and *Gjc1* (*Cx45*) in adult cardiac fibroblasts at 3 days after LAD-O or sham procedure. (D-G') HCF cultured in low or high density and stained for CX43 (green, D-E') and CX45 (green, F-G'). (H-I) Co-cultured iCMs (marked by TITIN-GFP, green) and HCFs stained for CX43 (red, H) or CX45 (red, I), with αSMA (grey). Arrowheads in H, gap junctions between iCMs. Arrows in H and I, gap junctions between iCM and HCF. (J-N) iCMs (J, K) and HCF (L, M) stained for $Fe^{2+}$ (red) after DMSO (J, L) or erastin (K, M) treatment. Fold change of $Fe^{2+}$ fluorescent intensity after erastin treatment in iCM and HCF quantified in N. Nuclei stained with DAPI (blue). Error bars indicate SD. **, p<0.01. Scale bar, 25 μm (A, B, D', E', F', G', H, I), 100 μm (D, E, F, G, J-M).

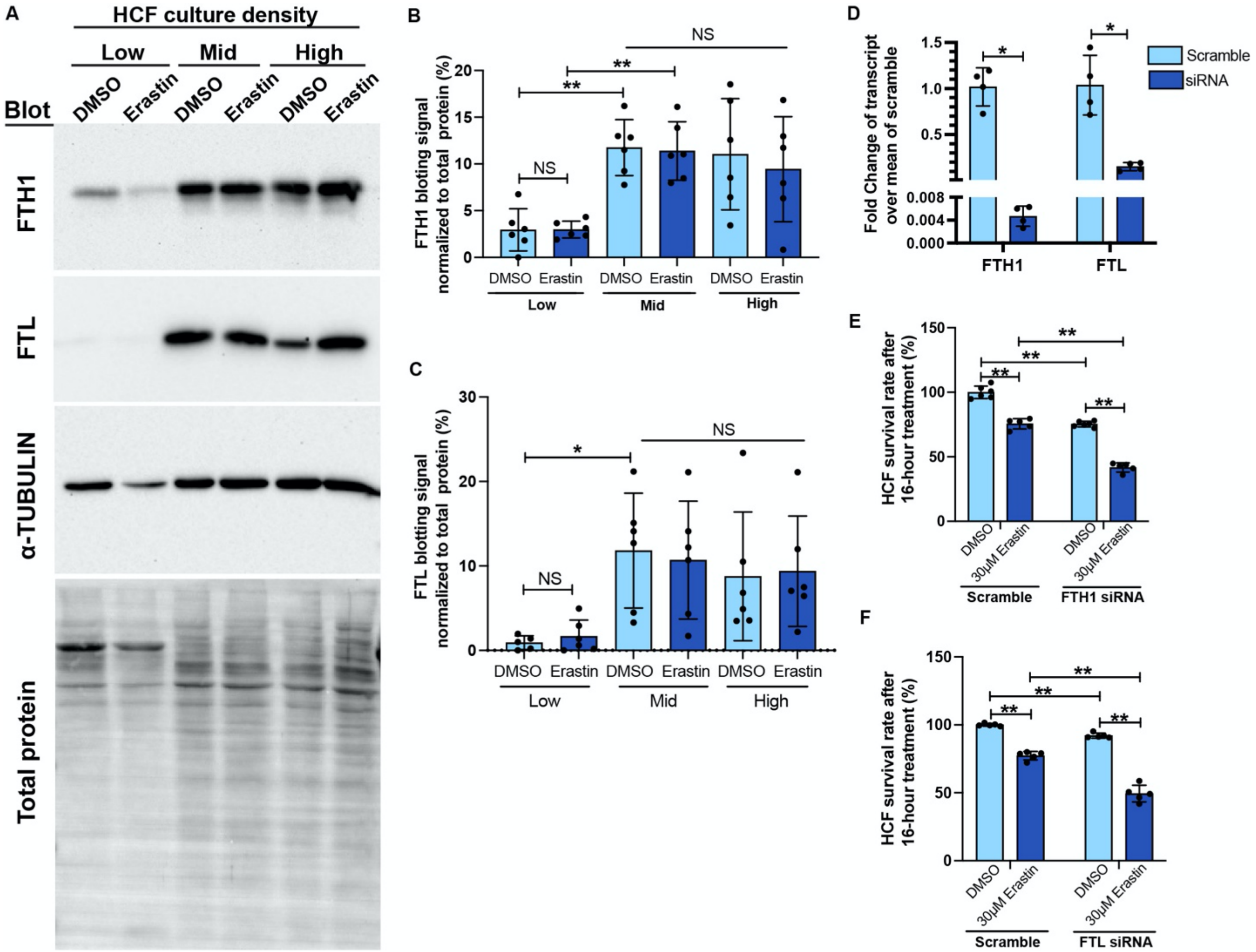


**Figure S5. Cardiac fibroblasts increase Ferritin expression to resist ferroptosis, related to Figure 5.** (A-C) Western blot of FTH1, FTL and α-TUBULIN in HCFs cultured at low, mid and high density, with DMSO or erastin treatment. Target band signal intensity quantified in B (FTH1) and C (FTL). (D) qPCR shows the knockdown of *FTH1* and *FTL* with siRNA in HCFs. (E, F) Survival rate of HCFs after erastin (30 μM) treatment, with *FTH1* (E) or *FTL* (F) knockdown compared to scramble siRNA. Error bars indicate SD. *, p<0.05. **, p<0.01. NS, not significant.

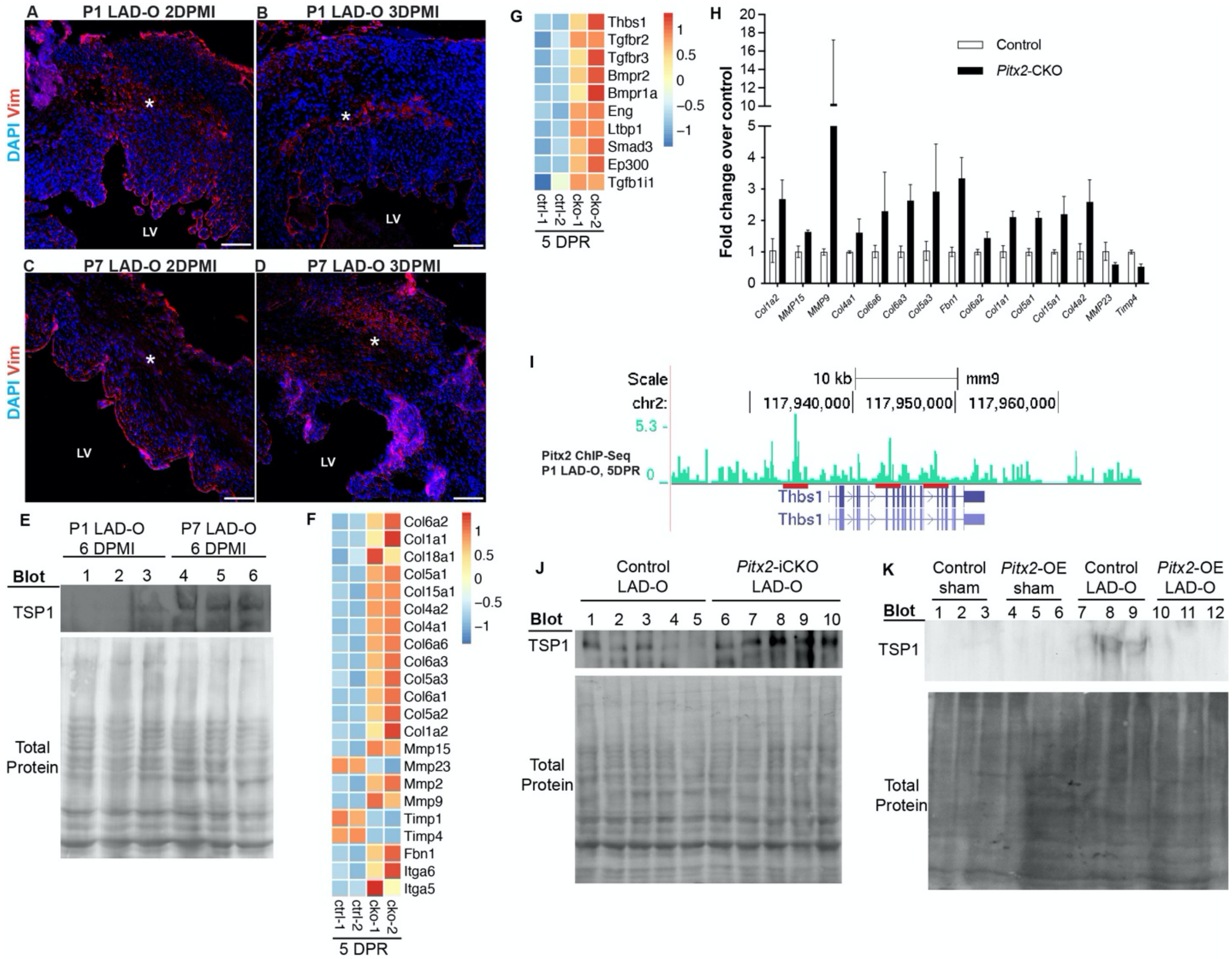


**Figure S6. Pitx2 regulates fibrotic gene expression in injured myocardium, related to Figure 6.** (A-D) Mouse heart tissue stained for Vim (red) at 2 or 3 days after P1 (A, B) or P7 (C, D) LAD-O. Asterisk, infarct zone. (E) Western blot of Tsp1 in cardiac ventricles at 6 days after P1 or P7 LAD-O. (F, G) Heatmap of fibrosis-relevant genes in control (*Pitx2$^{f/f}$*) and *Pitx2*-CKO (*MCK$^{cre}$;Pitx2$^{ff}$*) ventricles at 5 days after P1 apex resection. (H) qPCR validation of genes in F and G, $p<0.05$ for all targets. (I) ChIP-Seq shows Pitx2-binding region (red bars) at *Thbs1* locus in regenerative neonatal ventricles. (J) Western blot of Tsp1 in control and *Pitx2*-iCKO ventricles at 3 days after P1 LAD-O. (K) Western blot of Tsp1 in control and *Pitx2*-OE ventricles at 3 days after P7 LAD-O. LV, left ventricle. Nuclei stained with DAPI (blue). Error bars indicate SD. Scale bar, 75 μm (A-D).
